# Deconvolving the structural heterogeneity of alpha-Synuclein in *vitro* and *in situ*

**DOI:** 10.64898/2026.05.23.727349

**Authors:** Liliana Malinovska, Alexandra Malinovska, Yuehan Feng, Lynn Verbeke, Pratibha Kumari, Jose Camino, Valentina Cappelletti, Sonja Kroschwald, Elisa Dultz, Meltem Tatli, Walther J. Haenseler, Tetiana Serdiuk, Alexandra Estermann, Henning Stahlberg, Sally A. Cowley, Nunilo Cremades, Roland Riek, Lukas Reiter, Natalie de Souza, Paola Picotti

## Abstract

The structural states of proteins in cells and tissues provide important insight into their functional states, but studying protein structures *in situ* remains challenging. Furthermore, a single protein can adopt multiple conformations in cells, which typically cannot be assessed by most structural approaches. Here we developed a novel approach, based on structural proteomics fingerprints, for the quantitative analysis of the distribution of structural states of a protein *in vitro* and *in situ*. We applied it to the Parkinson’s disease hallmark protein alpha-synuclein (aSyn), for which various structural states (disordered, helical, oligomeric and amyloid fibrillar, among others) have been characterized *in vitro*, but for which the *in vivo* structural states remain hotly debated. We measured structure-specific proteolytic fingerprints from well-characterized aSyn *in vitro* conformations and used them to quantitatively determine the aSyn conformational composition in samples of interest. We first benchmarked our approach using ground truth datasets of known composition and showed that, during *in vitro* amyloid fibril formation, we could simultaneously detect a time-dependent decrease in disordered monomeric aSyn, an increase in β-sheet-rich oligomers, and a delayed rise in amyloid fibrils. We then applied the method to complex, biologically relevant samples. In a *S. cerevisiae* aSyn overex-pression model, aSyn was predominantly helical, with an increased helical fraction accompanying its relocalization from the plasma membrane to cytosolic lipid droplets. This shift was linked to proteome-wide changes in lipid droplet homeostasis and fatty acid and ergosterol metabolism, underscoring the role of lipid metabolism and droplet formation in aSyn biology. Importantly, we also detected helical aSyn in human iPSC-derived cortical neurons, supporting the physiological relevance of this conformation. Finally, neurons differentiated from PD patient-derived iPSCs showed elevated levels of β-sheet-rich aSyn compared to wild-type cells. Our approach allowed the *in situ* identification and quantification of different structural states of aSyn directly in cell lysates. Since several proteins can adopt multiple, functionally-relevant conformations in cells, our approach should be broadly applicable to *in situ,* quantitative structural and functional studies of proteins.

## Introduction

The structure of a protein dictates its functional state, and biological cues can alter protein structure^1^. Accordingly, aberrant protein structures can compromise protein function, resulting in disease^2^. For example, amyloidogenic proteins can assemble in beta-sheet rich amyloid fibrils, which are found in intra- and extracellular protein deposits in neurodegenerative diseases^3^. Many proteins are likely to exist in more than one structural conformation within cells and tissues. Thus, information on the collective of structural states of proteins can provide insight into the functional state of a system.

To advance in-cell structural biology, we have developed a new approach that enables analysis of the distribution of protein structural states *in vitro* and *in situ* (*i.e*., in cell lysates). It combines the capability of structural proteomics based on limited-proteolysis-coupled mass spectrometry (LiP-MS) to analyze protein structures directly from complex biological samples^4–8^, information from *in vitro* high-resolution structures, and the computational deconvolution of LiP-MS patterns from heterogeneous samples. Conformation-specific digestion patterns are first obtained from well characterized *in vitro* structural preparations. These are then compared to digestion patterns of the same proteins from potentially heterogeneous samples of interest, such as complex proteomes from cells or tissues. The contributions to the structural heterogeneity of a protein are determined using a multivariate mixing model.

We tested this approach using the protein alpha-synuclein (aSyn), which plays a key role in a set of neurodegenerative diseases called synucleinopathies^9–11^. In these diseases, which include Parkinson’s disease (PD), aSyn forms inclusions in dopaminergic and non-dopaminergic neurons as well as glial cells. A major challenge for elucidating the precise role of aSyn in neurodegenerative disease is its remarkable structural plasticity: It has been described as a structural chameleon^12^. *In vitro*, aSyn can adopt a variety of different structural conformations, some of which are interconvertible. In its monomeric form, aSyn is mostly disordered^13–15^. Upon binding to lipid vesicles, the monomeric form adopts a helical structure^16–19^. Upon prolonged incubation, monomeric aSyn aggregates into a beta-sheet rich amyloid fibril^14,20–23^. The formation of amyloid fibrils is preceded by the formation of unstructured and beta sheet-rich oligomeric species and protofibrils^24–27^. The beta sheet-rich oligomeric forms can cause membrane disruption and are associated with cellular toxicity^28–30^. Crucially, the structure that aSyn adopts *in situ*, within the cellular and tissue context, is still very controversial. In addition, the cellular and molecular consequences of aSyn pathology are not fully understood.

Given its plasticity, aSyn may exist in multiple, possibly interconvertible structures within the cellular and tissue environment, and the distribution of these structures may further be linked to the cellular state. Studying the functional role of aSyn in health and disease would be much aided by quantitative deconvolution of its *in situ* structural states. Here we applied our novel approach to analyse structural distributions of alpha-synuclein, both *in vitro* and in biologically and biomedically relevant cell lysates. We demonstrated using ground truth samples that the approach differentiated various structural states of aSyn both *in vitro* and in a complex background. We applied it to characterize structural transitions during amyloid fibril formation, and to determine the structural composition of aSyn in a yeast overexpression system and in neurons differentiated from patient-derived induced pluripotent stem cells (iPSCs). Notably, our data show experimental evidence for a helical aSyn structure within lysates of human iPSC-derived neurons and yeast, to our knowledge for the first time. In yeast, we found an association between the localization of aSyn to lipid droplets, formation of the helical conformation and concomitant alterations in the lipid homeostasis and LD biogenesis pathways. In neurons derived from PD patients, the aSyn helical fraction decreased with concomitant increase in beta sheet-rich structures and the aSyn structural composition was significantly different between control- and PD- derived neurons. Our deconvolution approach should be applicable also to other amyloidogenic disease-causing proteins with known structural plasticity (*e.g.*, TDP-43 in ALS or beta-amyloid in AD), and indeed to any protein that populates more than one structural state within the cell.

## Results

### Proteolytic fingerprints identify different structural conformations of aSyn

LiP-MS generates structure-specific proteolytic fingerprints of proteins even within complex biological samples. Thus, we speculated that comparing LiP patterns of aSyn structures generated *in vitro* and in cell or tissue lysates could be used to shed light on *in situ* aSyn structures and their heterogeneity. We prepared *in vitro* five well-characterized aSyn structures that are hypothesized to be present in biological samples: the unstructured monomer, a helical conformation induced by binding to small unilamellar vesicles (SUV)^31^, the kinetically trapped analogue of a non-toxic, predominantly unstructured oligomer (type-A*), and of a toxic, beta-sheet rich oligomer (type-B*)^25,27^, and amyloid fibrils^21^ (Table 1 and Fig. 1A). To account for variations that occur during sample preparation, we used three independent preparations of each aSyn conformation. Since amyloid fibril formation is known to be particularly variable^21,32–35^, we included amyloid fibrils generated in different laboratories starting from independent preparations of purified aSyn. To assess the structural effects of aSyn N-terminal acetylation, known to occur in mammalian cells^36^, we included acetylated and non-acetylated versions of the disordered monomer and amyloid fibrils. For all other structures, we used only the acetylated versions.

**Figure 1:**
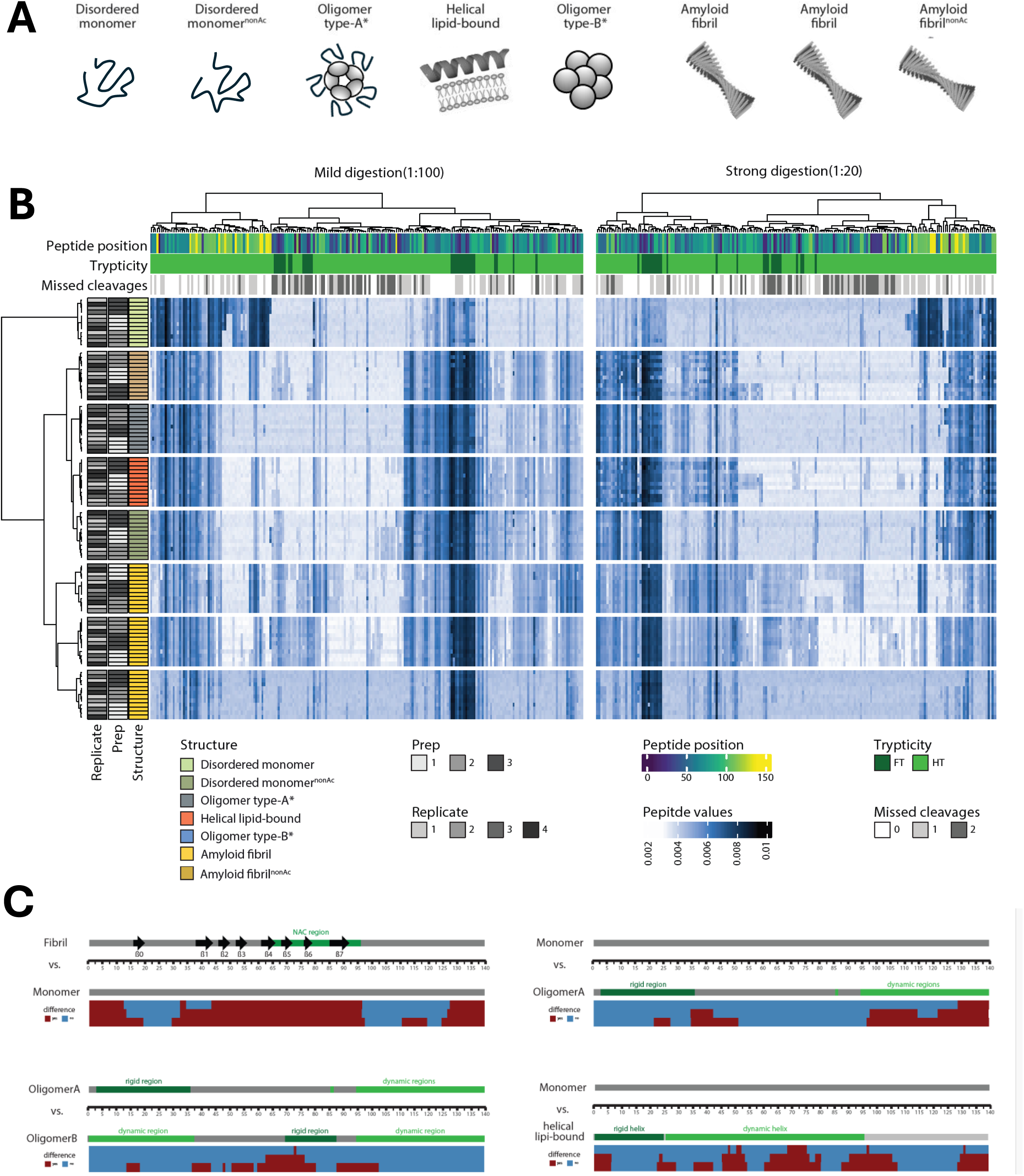
Structure-specific fingerprint library. **(A)** Schematic representation of structures included in the fingerprint library: disordered monomer (acetylated or non-acetylated); a kinetically trapped, non-toxic, predominantly unstructured oligomer (type-A*); a helical conformation induced by binding to small unilamellar vesicles (helical lipid-bound); a kinetically trapped, toxic, beta-sheet rich oligomer (type-B*); and amyloid fibrils from three different sources (two acetylated and one non-acetylated). **(B)** Unsupervised hierarchical clustering analysis of the structural fingerprints of the conformational samples. The transformed peptide intensities are visualized as feature values using a color gradient (white, low; dark blue, high). Row dendrogram: Different structural conformations described in A) are indicated by color. Individual preparations and technical replicates are indicated by shade of gray. Column dendrogram: Amount of missed cleavages are indicated by shades of gray, trypticity is indicated by color (dark green, fully tryptic; light green, half tryptic). The position of the center of the peptide is indicated by the viridis color scale. Distances are visualized in the dendrograms. **(C)** Pairwise comparison of the structure-specific digestion pattern of the reference structures (amyloid fibril vs. disordered monomer, disordered monomer vs. Oligomer type-A*, Oligomer type-A* vs. Oligomer type-b*, disordered monomer vs. helical lipid-bound). Structural features of the two conformations being compared are visualized along the sequence of aSyn. The results of the comparison shown in Fig. S2 is simplified and shown as binary outcome, visualized as a stacked histogram (red, structural difference; blue, no structural difference).

**Table 1:** Overview of purification and preparation protocols for the different structural conformations used in this study.

| Structural conformation | Purification protocol | Preparation protocol | Generated by |
| --- | --- | --- | --- |
| Disordered monomer, acetylated | 25 | 6 | Cremades Lab, University of Zaragoza |
| Disordered monomer, non-acetylated | 109,110 | 6 | Picotti Lab, ETH Zurich |
| Oligomer type-A* | 25 | 25 | Cremades Lab, University of Zaragoza |
| Helical lipid-bound | 25 | 31 | Cremades Lab, University of Zaragoza |
| Oligomer type-B* | 25 | 25,27 | Cremades Lab, University of Zaragoza |
| **Amyloid fibril,**<br>**Acetylated** | 108 | 6 | Picotti Lab/Riek Lab,<br>ETH Zurich |
| **Amyloid fibril,**<br>**non-acetylated** | 25 108,109 | 108 | Picotti Lab, ETH Zurich<br>Cremades Lab, Uni-<br>versity of Zaragoza |

We asked whether LiP-MS analysis of these five *in vitro* aSyn structures resulted in proteolytic fingerprints that were sufficiently different to allow detection and quantification of the structures in heterogenous samples. To generate LiP fingerprints, we briefly treated the samples with the non-specific protease proteinase K (PK) under either mild (1:100 enzyme:substrate, E:S) or strong (1:20 E:S) digestion conditions, followed by standard sample preparation and analysis of the resulting peptides by data-independent acquisition (DIA) mass spectrometry. Overall, we identified 460 proteolytic fragments (peptides); by excluding peptides below the detection limit in some replicates (Fig. S1A), we built fingerprints with a total of 406 peptide features (Fig. 1B) (210 peptides from the mild digestion condition and 196 peptides from the strong digestion condition) that map over the whole sequence of the protein (Fig. S1B).

We then assessed the discriminatory power of the different proteolytic fingerprints. Unsupervised hierarchical clustering showed a clear separation of fingerprints of the different conformations (Fig.1B). For all conformations, the independent preparations clustered together, indicating minimal differences in the sample preparation process. The amyloid fibril samples clustered as a separate branch, in which individual preparations were grouped predominantly based on the source (Table 1), rather than the preparation. Also, the acetylated monomer clustered as a separate branch, despite appearing minimally different in secondary structure from the non-acetylated form based on CD spectra^37^. Technical replicates were evenly distributed, indicating that there were no batch effects during the mass spectrometry acquisition step. The ability of proteolytic fingerprints to separate the six different structures was further confirmed using multiple data-dimensionality reduction techniques (Fig. S1C-E). These results demonstrate that proteolytic fingerprints generated by LiP-MS on defined *in vitro* samples are sufficiently distinct to differentiate the monomeric, helical, oligomeric (type-A* and type-B*) and amyloid conformations of aSyn.

Analysis of the proteolytic resistance of each aSyn conformation^38^, calculated as the ratio of peptide intensity in the LiP versus the trypsin-only condition (Methods) showed full proteolytic resistance (*i.e*., no statistical evidence for a difference in LiP and trypsin-only peptide intensities) between residues 13-96 for aSyn fibrils (Fig S1F), in agreement with the known rigid and protease inaccessible amyloid core of the fibril. In contrast, we observed a low proteolytic resistance for the monomer and the Oligomer type-A*, again consistent with findings from previous work^25^. The Oligomer type-B* and the helical lipid-bound conformations showed higher proteolytic resistance than the aSyn monomer, suggesting the presence of folded regions as previously reported^25,31^.

We further exploited the rich information in the complete structural fingerprints (consisting of 406 peptides) in pairwise comparisons of the five aSyn conformations and mapped the resulting structural differences along the aSyn sequence. We recapitulated many known structural features but also identified novel regions of structural difference between conformations (Fig. 1C, Fig. S2). Comparison of the monomer and fibrils showed differences across the NAC region (61-95) (Fig 1C), as expected for the fibrils we used in this study, which were prepared under conditions that would yield fibril polymorph 2^39^ (Figure S2B). Interestingly, we also observed differences between monomer and fibril in the N-terminus of the protein (1-13), the N-terminally adjacent region (30-61), and the C-terminus (125-140). Oligomer type-A* is generated by incubation of aSyn with EGCG^25,40^. Comparison to monomer showed a strong difference in the C-terminal part of aSyn (97-140) and a somewhat weaker difference in the region 35-50, consistent with our recent study showing that EGCG induced long range structural changes involving both aSyn termini^41^. Comparison of the two oligomeric forms, by contrast, showed differences in the NAC region and in the N-terminus, as expected from a previous report^25^. In the comparison of the monomer to the helical lipid-bound conformation, we observed an intriguing repetitive pattern of structural differences along the sequence. These included differences in the N-terminal region, consistent with the known N-terminal rigid helix anchored to the membrane, and in the known dynamic helix that determines the affinity of membrane binding^31^.

In summary, we have generated a library of LiP-MS based structural fingerprints of five conformations of aSyn that can discriminate between conformations, provides insight into the structural differences between them, and aligns with existing structural knowledge.

### A prediction model to assess conformational composition

Next, we used the five structural fingerprints of aSyn in a multivariate mixing model to determine the identity and composition of aSyn within an unknown sample. Here, the average fingerprints of the pure structures are defined as end members and the structure in an unknown sample is considered to be a mixture of these end members (Fig. 2A). This approach was first applied in ecological studies^42^ and was successfully adapted to use in other fields^43,44^.

**Figure 2:**
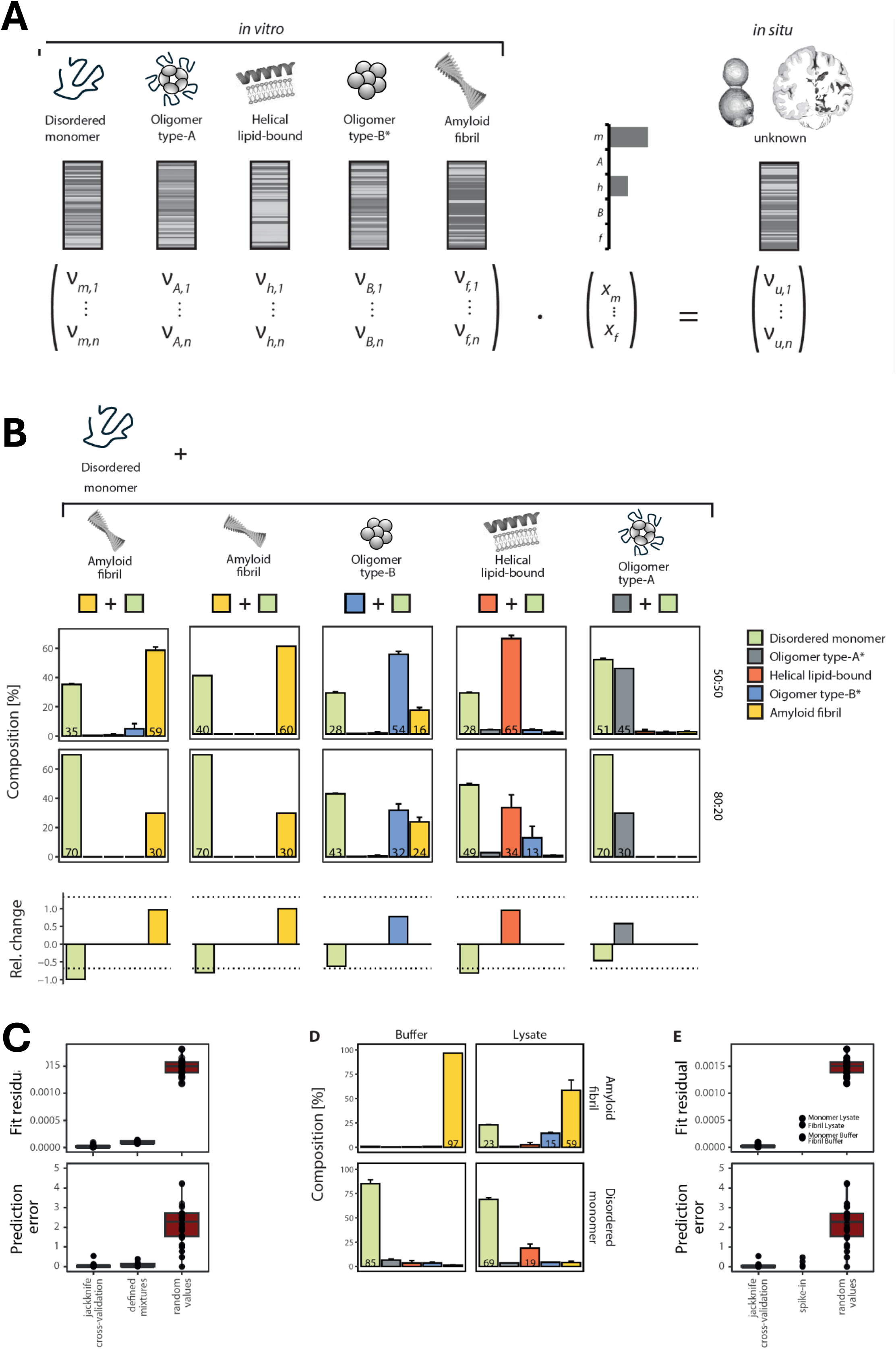
Multivariate mixing model for the prediction of conformational compositions. **(A)** Schematic representation of the prediction model. The peptide fingerprints of the in vitro structures are defined as end members in a multivariate mixing model and the composition of the unknown in situ sample is considered a mixture of those and determined by solving a linear inverse system. **(B)** Upper panel: Schematic representation of the test dataset. Acetylated disordered monomer was mixed with each of the other forms in 50:50 and 80:20 ratios. Middle panel: Results of the composition prediction for the 50:50 mixtures and 80:20 mixtures. The contribution of each structure is plotted as the fraction of the total (in [%]), and the identities of the predicted structures are indicated by color (disor-dered monomer, green; oligomer type-A*, grey; helical lipid-bound, orange; oligomer type-B*, blue; amyloid fibril, yellow). Errors are displayed as ranges and represent 95% confidence intervals of the Bayesian resampling. Lower panel: The relative changes of the two mixing partners in the comparison of the two mixing ratios. The fold changes are displayed in a bar plot, dotted lines represent the ex-pected changes. **(C)** Fit residuals (upper) and prediction error (lower) for the prediction in the mixture dataset described in B) (blue), jackknife cross-validation prediction (green), or prediction using random values (red). **(D)** Composition prediction for fibril and disordered monomer spiked into buffer (left) or cell lysate (right). The contribution of each structure is plotted as fraction of the total (in [%]) with identity of the predicted structures represented by color as in B. Errors are displayed as ranges and represent 95% confidence intervals of the Bayesian resampling. **(E)** Fit residuals (upper) and prediction error (lower) for the prediction in D (spike-in), jackknife cross-validation prediction (green), or prediction using random values (red).

To assess the suitability of the multivariate mixing model to predict each structural conformation, we used the structural fingerprints as a training dataset to fit a jackknife cross-validated prediction model and compared it to predictions using random values. We tested the performance of different types of fingerprints (Methods), all of which showed good performance in the cross-validation. In all cases, the model predicted the correct conformation as the dominant one (Fig. S3A), with fit residuals and prediction errors significantly lower than for predictions using random values (Fig. S3B). We selected one of these fingerprints, the robust fingerprint, for use in our prediction model since it showed the most even prediction of all conformations in a control where all peptides were assigned random values (Fig. S3C). Peptides of the robust fingerprint covered most of the sequence of aSyn well, except the N-terminal region of the protein, where it covered only residues 22-32 (Figure S3D).

Next, we assessed the performance of the model in predicting structural compositions of heterologous samples. We used a test dataset comprising predefined mixtures of the conformations used in the training dataset. We mixed the aSyn disordered monomer with each of the other aSyn conformations at two ratios (50:50 and 80:20; Fig. 2B). To avoid interactions between the different structures during sample preparation that could alter the known composition of a sample (*e.g*., conversion of the monomer into oligomers), we mixed the different conformations at the peptide level, after they had been processed by LiP-MS. Encouragingly, the model correctly predicted the two expected components as most abundant in all samples (Fig. 2B). For the mixtures containing only oligomer type-B*, the model predicted also a contribution from the amyloid fibril, consistent with our observations in the cross-validation tests (Fig S3A) and possibly due to their shared beta-sheet architecture^26,27^. A minor misprediction of the oligomer type-B* conformation in a mixture containing the helical form suggests that, only at low amounts, the helical lipid-bound conformation may not be easily distinguished from oligomer Type-B*. Whereas our approach did not recover the exact mixing ratios, the relative changes upon comparison of both ratios matched the expected ones very closely. Importantly, the fit residuals and prediction errors were lower than for the prediction of a single conformation using random values and only slightly higher than in the cross-validation of the training data (Fig. 2C). This suggests that the model can be used to determine the conformational components of a sample that contains aSyn.

Then, we assessed how prediction performance is influenced by a complex cellular background. We applied the prediction model to a validation dataset of samples where *in vitro* generated amyloid fibrils or disordered monomer were spiked either into buffer or into a cell lysate. Our model correctly predicted the expected structural form as the major component in all cases (Fig.2D). The fit residuals were comparable to those of the defined mixtures for the spike-in samples measured in buffer (Fig. 2E). For the samples measured in lysate, we observed higher fit residuals as well as prediction of small amounts of unexpected structures. This might indicate that aSyn interacts with lysate components and/or adopt additional conformations in this complex background; for instance, spiked-in monomer may interact with lipids in the lysate and take on the helical conformation. However, in all cases, the fit residuals were lower than those for random values (Fig. 2E) and the prediction error was comparable to that of the defined mixtures (Figure 2C, E).

Taken together, these results show that our approach can be used to predict the main structural states of a protein in both purified and complex biological samples.

### Prediction of conformational composition over an aggregation time course

We next applied our model to low-complexity samples of unknown composition. We sampled aSyn over time under conditions that result in aggregation of disordered aSyn monomers to amyloid fibrils, adding aSyn seeds at time zero (t0) to promote more rapid and coherent aggregation. Thioflavin T (ThT) fluorescence^45,46^ showed amyloid fibril formation starting from day 4 (Fig. 3A), while our model indicated a small increase in amyloid content starting already from day 2 (Fig. 3B), which may reflect differences in sensitivity and dynamic range of the two approaches. Nevertheless, our approach showed an increase in amyloid and a decrease in the disordered monomer over time, as expected. It further showed an initial increase in type-B* oligomers, followed by their decrease as amyloid fibrils accumulate.

**Figure 3:**
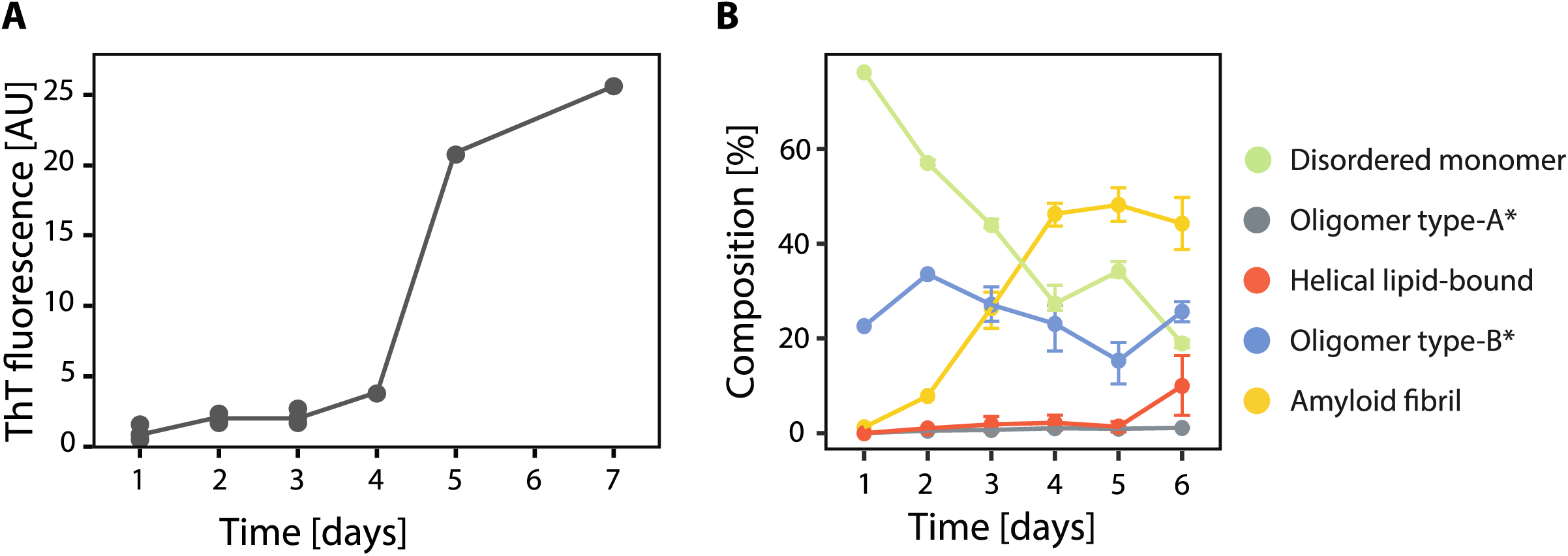
Prediction of aSyn conformation over time under aggregation-promoting conditions. **(A)** Formation of amyloid fibrils over the course of 7 days monitored with ThT fluorescence. Replicates (n=3) are visualized as dots; the mean value is visualized as line. **(B)** Predictions of conformational composition over time as fraction of the total (in [%]). The identities of the predicted structures are represented by color as indicated. Errors are displayed as ranges and represent 95% confidence inter-vals of the Bayesian resampling.

The low fit residuals for these predictions indicate high confidence (Fig. S3E), and indeed our predictions agree with previous observations that disordered monomers assemble first into transient, disordered type-A* oligomers, which are converted into beta-sheet containing type-B oligomers and ultimately into amyloid fibrils (Fig. 3B)^26^. The addition of seeds to the aggregation assay likely contributed to identification of the oligomer type-B* at the earliest time point. The predicted increase in oligomer type-B* at day 6 may be due to the known tendency of amyloid fibrils to disassemble into this form in monomer-free solutions.^26,27^ Interestingly, our model also suggested the formation of small amounts of a helical conformation at day 6, in agreement with a recent study showing that helix-rich intermediates can form during aSyn aggregation^47^. Overall, these results show that the model can predict the composition of an unknown *in vitro* sample of low complexity.

### aSyn adopts a helical conformation and colocalizes with lipid droplets in yeast

We applied our approach to analyze aSyn structural heterogeneity in complex cellular samples. We focused first on yeast overexpressing aSyn, a broadly used model for studying aSyn toxicity^48^, in which aSyn localizes first to the plasma membrane and then translocates to cytosolic accumulations, comprising membranous vesicular clusters^49^. We recapitulated previous toxicity data of this model using a two-fold serial dilution assay^50^ (Fig. S4A). Using a fluorescently tagged aSyn construct in an otherwise identical strain, we observed predominantly plasma membrane localization three hours post-induction (Fig.4A), whereas at 12 hours post-induction, only 32% cells showed aSyn plasma membrane localization, 36% of the cells had small cytosolic foci, and 32% had large cytosolic accumulations (Fig. 4B, Fig. S4B).

**Figure 4:**
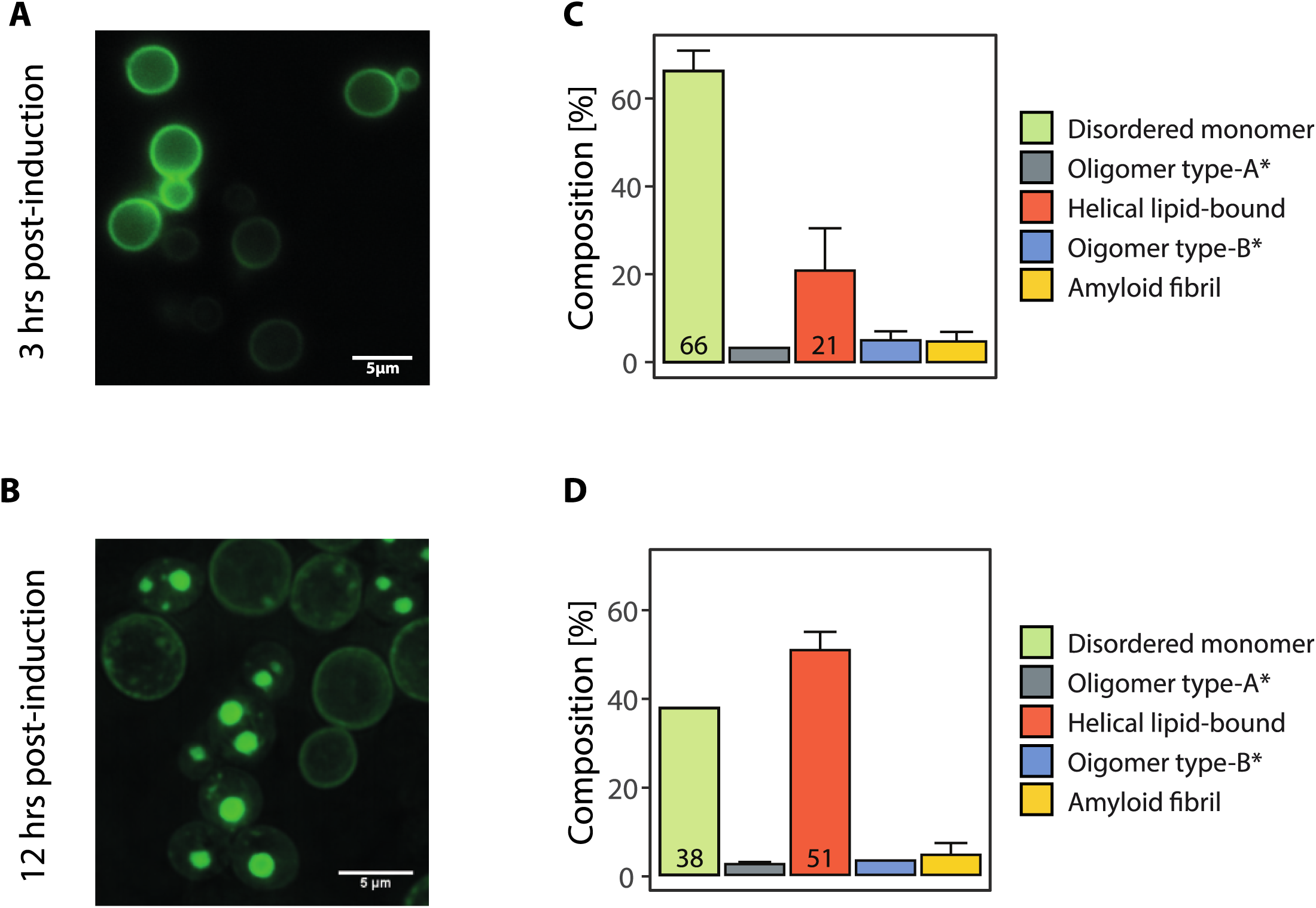
Prediction of aSyn conformations in a yeast overexpression system. (A, **B)** Fluorescent light microscopy characterization of *S. cerevisiae* that express GFP-tagged aSyn at **A)** 3 h post-induction and **B)** 12 h post-induction. Scale bar 5 µm. **(C, D)** Composition prediction of aSyn in **C)** *S. cerevisiae* lysates aSyn at 3 h post-induction and **D)** *S. cerevisiae* lysates aSyn at 12 h post-induction. The identities of the predicted structures are represented by color, as indicated. Errors are displayed as ranges and represent 95% confidence intervals of the Bayesian resampling.

Our analysis identified the disordered monomer and helical conformation within the yeast lysate at both time points (Fig. 4C, D). As we saw for the spikeins to cell lysate, the fit residuals for these predictions were higher than for samples in buffer, likely reflecting background complexity, but substantially lower than those obtained with random values (Fig. S3E). Interestingly, the fraction of helical aSyn increased at 12 hours post-induction (Fig. 4D), concomitant with the translocation of aSyn-GFP to cytoplasmic inclusions (Fig. 4B). Staining with Nile Red showed that these aSyn-GFP inclusions co-localized well with lipid droplets (Fig. S4C) as has been previously observed^51–53^. Induction of either untagged or GFP-tagged aSyn also resulted in an overall increase in the number of LDs (Fig. S4D) as expected^54^. This suggests the acquisition of a helical structure by aSyn upon association with membranes and the accumulation of this conformation, increasingly bound to LD membranes, over time. Intriguingly, comparison of the aSyn structural fingerprints between early and late time points showed a repetitive pattern of structural differences (Fig. S4E), which could be consistent with the formation of increasing amounts of a helix.

Using fluorescence recovery after photobleaching (FRAP) data we further confirmed that aSyn-GFP foci formed at 12h are not amyloids, since they show a substantially higher median mobile fraction (40%) than that of the well-characterized amyloid-forming yeast prion PSI+ (20%) (Fig. S4F). These data and the very low amount of amyloid fibrils predicted by our model agree with previous studies in which no fibril formation could be detected in *S. cerevisiae*^49^. We also analyzed these cells using high resolution cryo-ET and observed no fibrillar structures co-localizing with GFP-positive foci (Fig. S4G).

The possible interaction between aSyn and LDs prompted us to make use of the proteome-wide LiP-MS data generated as part of this study to assess whether alterations in the biology of LDs or lipid metabolic pathways can be detected upon overexpression of aSyn. We previously showed that LiP-MS data can detect proteome-wide structural changes of proteins due to cataysis, allostery, altered protein-protein interactions, binding to other molecules, post-translational modification, aggregation and unfolding events, thereby capturing a variety of functional changes in a proteome, including regulation of metabolic pathways^7,8,55–57^ We therefore probed the dataset for global structural alterations and identified proteins that show structural as well as abundance changes at 12h post-induction in comparison to a non-induced sample. Functional enrichment analysis on this set of proteins using the Kyoto Encyclopedia of Genes and Genome (KEGG)^58^ metabolic pathways and the Gene Ontology (GO)^59^ cellular components showed an enrichment in lipid metabolism (Fig.5A, purple box), as well as in lipid droplets and the plasma membrane compartment (Fig.5B, purple box). We also observed an enrichment in proteins involved in carbohydrate and amino acid metabolism and in translation, most probably associated with the overexpression system (Fig.5A, B; blue boxes). We detected structural alterations in known interactors of aSyn (Fig. S5A) as well as in previously identified modulators of aSyn toxicity (Fig. S5B),^48,51–53^ and identified numerous structural changes in proteins associated with fatty acid biosynthesis, ergosterol synthesis,^60,61^ and lipid/lipid droplet biology (Fig. 5C)^62,63^, suggesting regulation of these pathways concomitant with the co-localization of aSyn with LDs.

**Figure 5:**
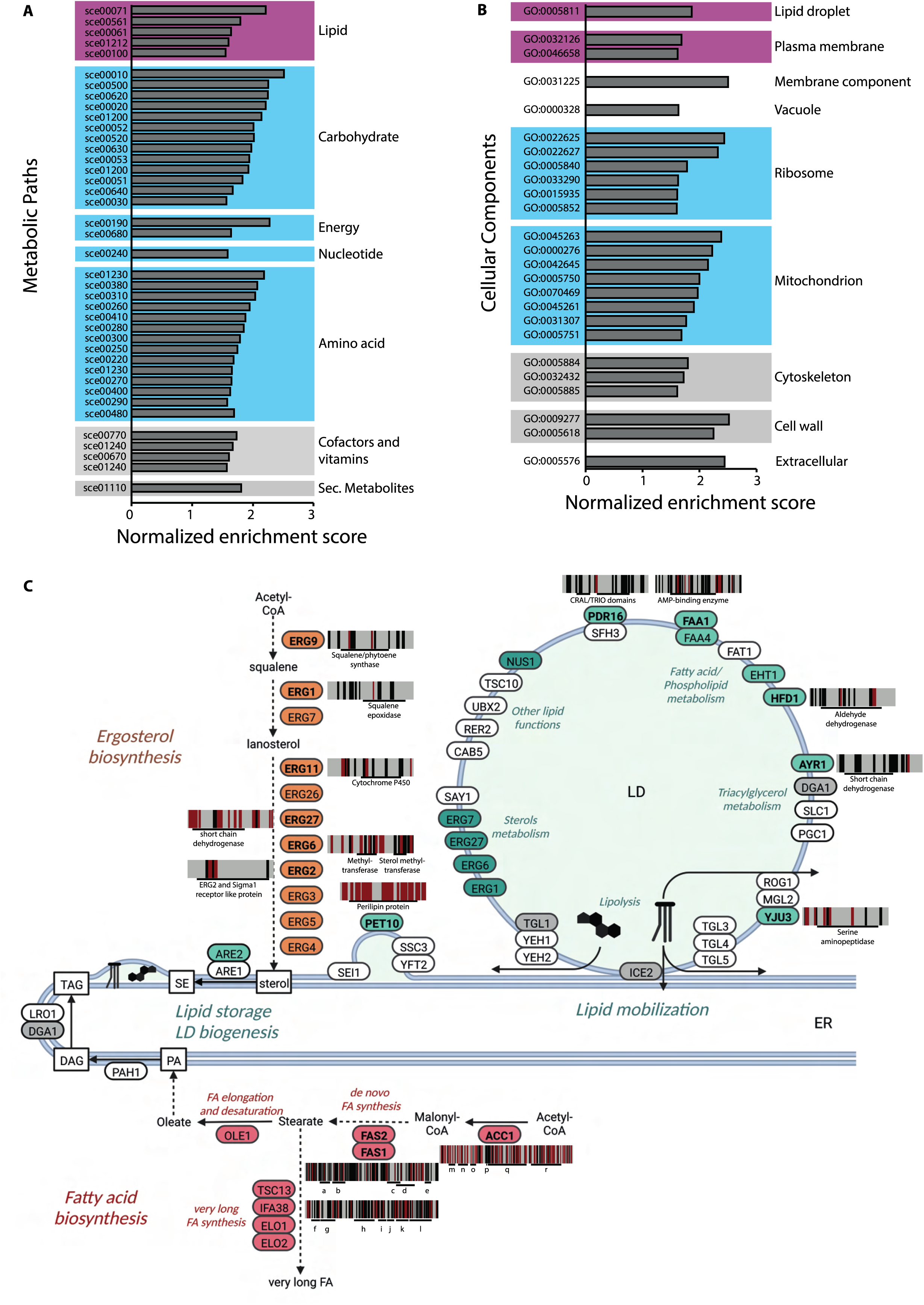
Global alterations upon aSyn overexpression in yeast. (A, **B)** Protein set enrichment analysis usin g **A)** Kyoto Encyclopedia of Genes and Genome (KEGG) metabolic pathways and **B)** the Gene Ontology (GO) cellular components. Proteins showing abundance changes and structural alterations under mild digestion conditions upon aSyn overexpression 12 hours post-induction are compared to a non-induced sample. Terms are grouped manually and plotted against the normalized enrichment score. Those associated with the overexpression system are highlighted in blue, groups related to lipid metabolism are highlighted in purple. sce00071, Fatty acid degradation; sce00561, Glycerolipid metabolism; sce00061, Fatty acid biosynthesis; sce01212, Fatty acid metabolism; sce00100, Steroid bio-synthesis; sce00010, Glycolysis / Gluconeogenesis; sce00500, Starch and sucrose metabolism; sce00620, Pyruvate metabolism; sce00020, Citrate cycle (TCA cycle); sce01200, Carbon metabolism; sce00052, Galactose metabolism; sce00520, Amino sugar and nucleotide sugar metabolism; sce00630, Glyoxylate and dicarboxylate metabolism; sce00053, Ascorbate and aldarate metabolism; sce01200, Carbon metabolism; sce00051, Fructose and mannose metabolism; sce00640, Propanoate metabo-lism; sce00030, Pentose phosphate pathway; sce00240, Pyrimidine metabolism; sce01230, Biosynthe-sis of amino acids; sce00380, Tryptophan metabolism; sce00310, Lysine degradation; sce00260, Gly-cine, serine and threonine metabolism; sce00410, beta-Alanine metabolism; sce00280, Valine, leucine and isoleucine degradation; sce00300, Lysine biosynthesis; sce00250, Alanine, aspartate and gluta-mate metabolism; sce00220, Arginine biosynthesis; sce01230, Biosynthesis of amino acids; sce00270, Cysteine and methionine metabolism; sce00400, Phenylalanine, tyrosine and tryptophan biosynthesis; sce00290, Valine, leucine and isoleucine biosynthesis; sce00770, Pantothenate and CoA biosynthesis; sce01240, Biosynthesis of cofactors; sce00670, One carbon pool by folate; sce01240, Biosynthesis of cofactors; sce00480, Glutathione metabolism; sce01110, Biosynthesis of secondary metabolites; GO:0005811, lipiddroplet; GO:0032126, eisosome; GO:0046658, anchored component of plasma membrane; GO:0031225, anchored component of membrane; GO:0000328, fungal-type vacuole lu-men; GO:0022625, cytosolic large ribosomal subunit; GO:0022627, cytosolic small ribosomal subunit; GO:0005840, ribosome; GO:0033290, eukaryotic 48S preinitiation complex; GO:0015935, small ribo-somal subunit; GO: 0005852,eukaryotic translation initiation factor 3 complex; GO:0045263, proton-transporting ATP synthase complex, coupling factor F(o); GO:0000276, mitochondrial proton-transporting ATP synthase complex, coupling factor F(o); GO:0042645, mitochondrial nucleoid; GO:0005750, mitochondrial respiratory chain complex III; GO: 0070469, respirasome; GO:0045261, proton-transporting ATP synthase complex, catalytic core F(1); GO:0031307, integral component of mitochondrial outer membrane; GO:0005751, mitochondrial respiratory chain complex IV; GO:0005884, actin filament; GO:0032432, actin fi lament bundle; GO:0005885, Arp2/3 protein com-plex; GO: 0009277, fungal-type cell wall; GO:0005618, cell wall; GO:0005576, extracellular region. **(C)** Schematic representation of lipid storage and lipid droplet (LD) biogenesis, lipid mobilization in LDs, fatty acid and ergosterol biosynthesis. Protein with a significant structural alteration (q-value <0.05) are highlighted in color (LD-associated proteins, turquoise; fatty acid biosynthesis, red; ergosterol bio-synthesis, orange). Proteins found in the data set, but with no structural alteration are highlighted in grey, proteins not identified in the data set in white.Proteins leading the enrichment (see Methods for more details) are highlighted bold. Peptides displaying a significant change in the mild digestion condi-tion are plotted along the sequence of the protein and highlighted in red. Peptide with no significant change are shown in black, positions not identified are shown in grey. Pfam protein domains are indi-cated below the protein sequence. Protein domains in FAS2: a, Fatty acid synthase subunit alpha Acyl carrier domain; b, Fatty acid synthase type I helical domain; c, Beta-ketoacyl synthase, N-terminal domain; d, Beta-ketoacyl synthase, C-terminal domain;’ e, 4’-phosphopantetheinyl transferase super-family. Protein domains in FAS1: f, N-terminal domain in fatty acid synthase subunit beta; g, Starter unit:ACP transacylase in aflatoxin biosynthesis; h, Domain of unknown function; i, Fatty acid synthase meander beta sheet domain; j, N-terminal half of MaoC dehydratase; k, MaoC like domain; l, Acyl transferase domain; m, Biotin carboxylase, N-terminaldomain; n, Carbamoyl-phosphate synthase L chain, ATP binding domain; o, Biotin carboxylase C-terminal domain; p,Biotin-requiring enzyme; q, Acetyl-CoA carboxy-lase, central region; r, Carboxyl transferase domain. TAG,triacylglycerol; DAG, diacylglycerol; PA, phosphatidic acid; ER, endoplasmic reticulum; SE, sterol ester

In summary, our quantitative *in situ* analysis of aSyn structural heterogeneity indicates that aSyn in the yeast overexpression system adopts a largely helical and likely lipid-associated conformation over time and that the fraction of this conformation increases concomitant with alterations in yeast lipid metabolism and the lipid droplet proteome.

### aSyn structural fingerprints identify a helical conformation in iPSC-derived neuronal lysates

To test our findings in a more physiologically relevant system, we next analyzed aSyn conformations in neurons differentiated from patient-derived iPS cells. We compared cell lines with wild type levels of aSyn derived from healthy controls (n=3 cell lines from 3 individuals) with cell lines with a triplication of the SNCA gene locus derived from a Parkinson’s disease patient (n=3 cell lines from a single individual). This genomic alteration causes a familial form of PD^64^ and results in 2-3 fold higher aSyn expression compared to the healthy control samples.

The aSyn full proteolytic fingerprint from control and PD-derived samples clearly separated in a PCA (Fig. 6A), indicating different structural features of aSyn in the two samples. In iPSC-derived neurons with wild type aSyn, we predicted the helical form as the major conformation, with a small contribution from amyloid structures. In cells with SNCA triplication, the contribution of the amyloid fibril was substantially increased, and we also predicted the presence of oligomer type-B* (Fig. 6B). The fit residuals for these predictions were lower than random, and higher than the residuals for sample in buffer, again likely reflecting background complexity (Figure. S3E). Taken together, our data suggest an increase in beta sheet-rich conformations in neurons differentiated from PD patient-derived iPSCs relative to wild type cells. Consistent with this, immunofluorescence microscopy with an antibody against aggregated aSyn showed more and larger cytosolic aSyn foci in triplication than in WT cells (Fig. 6C, D). Moreover, a recent study also reported an increase in beta sheet-rich content in aSyn triplication iPSC-derived spheroids using Fourier transform infrared microspectroscopy^65^.

**Figure 6:**
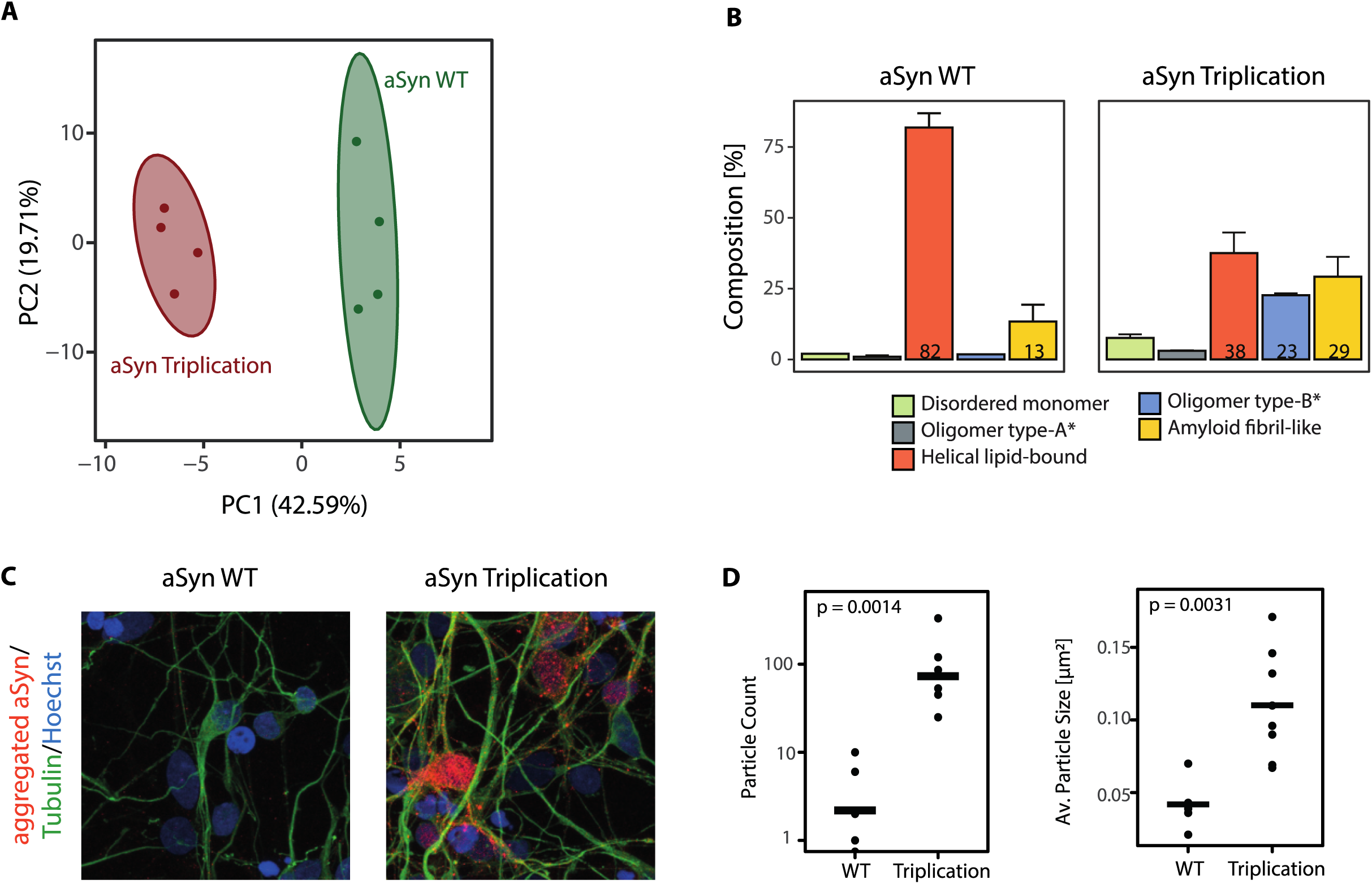
Composition analysis of aSyn in patient iPSC-derived cortical neurons. **(A)** Principal component analysis of the aSyn structural fingerprint from pooled lysates of iPSC-derived neurons with wild type aSyn (green; n=3 iPSC lines from 3 individuals) and aSyn triplication (red; n=3 iPSC lines from one PD individual). Each dot indicates an individual LiP-MS technical replicate. **(B)** Composition prediction in lysates as in A, with wild type aSyn (left) and aSyn triplication (right). The predicted structures are indicated by color as indicated. Errors are displayed as ranges and represent 95% confidence intervals of the Bayesian resampling. **(C)** Fluorescence images of iPSC-derived cortical neurons showing aSyn staining with an anti-aggregated aSyn antibody, tubulin staining, and nuclear staining with Hoechst. **(D)** Quantification of image data as in C in neurons derived from all lines used for the LiP analysis, showing aSyn particle number (left) and size (right). The individual points indicate images (n=7-8 images from 2-3 iPSC lines) and the horizontal line shows the mean.

Overall, our analysis predicts a predominantly helical conformation of aSyn in wild type iPSC-derived neurons, and a shift to beta sheet-rich conformations in PD patient-derived cells.

## Discussion

Many proteins, in particular proteins with intrinsically disordered domains, can adopt multiple conformations, such that a mechanistic understanding of protein function requires knowing which structures exist within cells or tissue. The structure that the key Parkinson’s disease protein aSyn adopts *in situ* remains very controversial. In-cell NMR studies of monomeric aSyn electroporated into mammalian cells have shown that the protein remains monomeric within cells and adopts a more compact structure than in buffer,^66^ and interacts transiently with chaperones^67^. Some studies have reported the presence of stable helical tetramers^68–70^, whereas others have provided evidence for a stable disordered monomeric form^15,39,66,71,72^. Similarly, the pathological conformation of aSyn within cellular deposits in patients is not clear. Early studies showed that aSyn forms fibrillar amyloid-like structures and these structures have long been thought to define Lewy Bodies (LB) in PD^73–75^. However, high-resolution EM and ET has more recently shown that these inclusions are rich also in membranous and organellar material, in some cases interspersed with fibrillar structures.^76^ Here we present the first quantitative analysis of the *in situ* conformational distribution of aSyn. Our study shows, to our knowledge for the first time, experimental evidence for a helical aSyn structure within yeast and human iPSC-derived neurons based on *in situ* measurement of endogenous aSyn structural properties in cell lysates.

Our data provide experimental evidence for recent predictions, based on deep mutational scanning, suggesting that a helical form of aSyn is relevant for its toxicity in yeast^77^. A helical aSyn conformation could be in agreement with various proposed helical variants, some of which have been shown to associate with membranes^25,31,78,79^. We note that, although the *in vitro* structure from which we derived the helical fingerprint is lipid-bound, our data do not directly demonstrate that the helical conformation we predict is bound to lipids *in situ*. Nevertheless, several observations are consistent with our predictions reflecting a lipid-bound helical structure. We showed in yeast that the fraction of helical aSyn increased concomitant with the formation of aSyn-positive cytoplasmic inclusions. These inclusions co-localized with lipid droplet (LD) markers, as we and others have also shown previously in yeast^80,81^ and mammalian cells^82,83^. aSyn overexpression also induces the formation of LDs in yeast^48,81^ and mammalian cells^82–84^, which is enhanced when the cells are treated with lipids. Further, there are important similarities between the membrane-targeting motif of aSyn and that of perilipins, proteins that bind the surface of lipid droplets^85,86^. aSyn undergoes a conformational change from intrinsically disordered to a largely helical structure upon binding to lipids *in vitro*. This is driven by a string of 7 imperfect 11-mer repeats that form amphipathic helical structures upon lipid interaction, and facilitate αSyn binding to a diverse array of lipid environments, including micelles, lipid vesicles, and membranes^87,88^. Perilipins contain the same 11-mer repeats as aSyn, and these are thought to be important in the regulation of lipid-protein interactions^85^. Taken together, it is therefore very likely that the *in situ* helical aSyn structure we detect represents aSyn bound to membranes, but this will need confirmation with orthogonal *in situ* methods.

The toxicity of aSyn in yeast overexpression models is linked to its membrane binding^89^, and our global analysis of structural changes suggests that this may at least in part involve a deregulation of LD biology. We observed that accumulation of helical aSyn structures and aSyn-positive LDs upon aSyn overexpression in yeast were accompanied by many protein structural changes, with an enrichment of LD-associated proteins^62,63^ among the set of structurally altered proteins. These could reflect direct interactions of aSyn with LD-associated proteins, changes in the curvature or rigidity of the droplet surface due to aSyn binding, or regulation of lipid metabolism enzymes associated to the droplets. Consistent with this, we recently showed that inhibition of the phosphatidate phosphatase Pah1, which controls LD formation rescues cells from aSyn toxicity^80^. In addition, several previously identified modulators of aSyn toxicity are closely connected to Pah1 in a functional interaction network and are likely to modulate this toxicity through the same lipid biosynthetic pathway^80^. Previous work has also shown that the increase in LDs following aSyn overexpression in yeast was associated with an increase of oleic acid triggered by upregulation of acetyl-CoA carboxylase (Acc1) and stearoyl-CoA desaturase (Ole1), two proteins we also find structurally altered^54^.

Beyond LDs, lipid dyshomeostasis in general is proposed to be part of the adaptive response to aSyn overexpression^53,54,80,84^. In agreement with this, we find several proteins within lipid metabolism, all of them previously implicated in aSyn toxicity,^53,84^ that change structure and are thus likely to be functionally altered upon aSyn overexpression e.g., both subunits of the fatty acid synthase (FAS1 and FAS2), proteins associated with very long chain fatty acid bio-synthesis, as well as components of the ergosterol biosynthesis pathway. Further analyses of the relationship between the fraction of aSyn that is helical, the extent of LD formation and cellular toxicity may clarify the role of the helical structure in the pathophysiology of aSyn

aSyn has also been linked to lipid metabolism in mammalian systems^51,80,81^. In mammalian cells, aSyn also localizes to LDs,^31,54^ overexpression of aSyn induces lipid droplet formation,^31,51,52,54^ and aSyn is linked to neuronal fate through interactions with neutral lipid metabolism^51^. In-cell FRET experiments in mammalian cells are also compatible with a helical conformation of vesicle-localized aSyn^83^. Interestingly, more than 15 PD- or parkinsonism-related genes are involved in membrane or lipid biology. One of them *DGKQ,* emerged as PD risk factor in independent GWAS studies^90,91^ and regulates the cellular content of DAGs, the precursors of LDs.^92^ Aggregates of aSyn associate with vesicular structures in a mammalian cell model of Lewy body biogenesis^54^. In PD patients, aSyn-rich Lewy bodies and Lewy neurites contain membranous material, including vesicular structures^76^. Both Lewy bodies and the inclusions observed in the cellular model also contain LDs^54^. Physiologically, aSyn has been linked to membrane-binding (reviewed in^84^), to the endolysosomal system (reviewed in^89^) and synaptic vesicle homeostasis (reviewed in^93^). A recent study showed that the docking of synaptic vesicles to the presynaptic membrane by aSyn depends on the lipid composition^94^, suggesting the relevance of aSyn lipid binding, and thus of the helical conformation detected in our study, under both physiological and pathological conditions.

Our method to assess the composition of aSyn structures in an *in situ* sample is based on a multivariate mixing model, an approach previously applied in ecological studies^42^. We used the fit residuals to assess the performance of the prediction model. Interestingly, the fit error was consistently lower for samples in buffer than in a complex background. This could suggest the presence of an as-yet unknown structure or one that is not included in the library; indeed a limitation of the approach is that the library does not include all possible structures of aSyn. For instance, aSyn has been reported to have multiple, interconvertible binding modes to lipids ^16–19,95^, as well as to form large lipoprotein particles ^96–98^ and undergo liquid-liquid phase separation ^99,100^. We further did not include in our *in vitro* library the stable helical tetramers that have been proposed to occur *in vivo*^68–70,101–104^ but were challenged by others^66,72,105^, because these structures require chemical cross-linking for stabilization and purification and are therefore incompatible with our LiP-MS workflow. Errors in the fit can also be caused by interactions of aSyn with proteins from the lysate such as chaperones^67^, which would alter protease accessibility and therefore affect our structural fingerprints, or originate from MS-associated technical sources where the background causes ion suppression effects that affect peptide intensities.

When characterizing the *in vit*ro generated structures, we detected minor differences in aSyn structures from different preparations of the same structure, even though we used only structures with reproducible preparation protocols^25,27,31^. In particular, we observed differences in aSyn amyloid fibrils according to the source of protein and the preparation. This is an important observation for both structural and functional studies of amyloids. Different structures of preformed fibrils (PFF) used in seeding and propagation studies (reviewed in^106^) could translate into different outcomes, impeding the comparison and interpretation of results. Interestingly, our data indicate that the non-acetylated aSyn monomer was structurally more similar to the oligomeric forms than to the acetylated monomer, while acetylation did not affect fibril structure. This observation is surprising and may hint at acetylation having more of an effect on the structure of disordered aSyn monomer than on the more structured fibril. We note as well that our LiP experiments were carried out at 25°C, and recent studies showed a temperature-dependent conformation switch of aSyn in neurons^107^. Future studies will assess the temperature-dependence of our predicted *in situ* structures.

Taken together, we have developed an approach to quantitatively determine the structural landscape of proteins within samples with unknown composition, limited solely by the number and type of well characterized structural conformations that can be included in a structural fingerprint reference library. Importantly, our approach can be used *in situ* to deconvolve the structural states of a protein of interest within a complex, physiologically relevant context and to assess the *in situ* occurrence of high-resolution structures from *in vitro* studies. The predictions of our approach were in good agreement with prior knowledge and have provided evidence for a helical, likely lipid-associated structure of aSyn within yeast cells and in iPSC-derived human neurons. Our approach should be applicable to any intrinsically disordered or amyloidogenic protein, or indeed any protein with multiple conformational states, and should yield insight both into which structural states of a protein are present in cells and tissues, and how changes in the distribution of these states upon perturbation, such as disease, may influence function.

## Supporting information

Malinovska et al Supplement

## Acknowledgements

We thank Senthil Kumar and Hilal Lashuel for insightful discussions on the centrifugation-based filtration protocol. We thank Ilaria Piazza for constructive discussions on the sample preparation and MS/MS acquisition strategy. We thank Lucie Kralickova for her support in the purification of aSyn. LM was supported by the long-term EMBO postdoctoral fellowship (ALTF 538-2016). WH was funded by the Oxford-McGill-Zurich Partnership in Neuroscience and UZH URPP AdaBD. P.P. was funded by the Peter Bockhoff Stiftung and the ETH Zurich foundation, Parkinson Schweiz, an EMPIRIS Foundation grant (2022-FS-353) and the Synapsis Foundation - Alzheimer Research Switzerland ARS.

## Data Availability

All proteomics data generated in this study will be available from PRIDE. Accession numbers will be provided before publication.

## Code Availability

Code is deposited at https://gitfront.io/r/PicottiGroup/GffTFNnJa2My/Deconvolving-the-struc-tural-heterogeneity-of-aSyn-in-vitro-and-in-situ/. All code central to this study will be available upon publication.

## Author contributions

L.M. designed and performed the experiments and supervised aspects of the computational pipeline, L.M. and A.M. designed and performed the bioinformatics analysis of the data, Y.F. and L.R. contributed to the project design, V.C., L.V. and L.R. contributed to the data analy-sis, P.K. and J.C. provided preparations of different aSyn conformations, S.K. performed FRAP experiments and analysis, E.D. performed LD staining and microscopy, MT performed cryo-ET, W.H. grew iPS cells and performed IF staining, T.S. contributed to aSyn fibril prepa-rations, A.E. contributed to experiments, S.A.C. provided iPS cell lines, H.S. provided re-sources and support for cryo-ET, L.M, A.M., N.d.S. and P.P. wrote the manuscript, N.d.S. contributed to supervision, P.P. conceived and supervised the project.

## Methods

### Protein purification and preparation of aSyn conformations

The different aSyn conformations were prepared in three independent preparations as described previously. Details on the expression of recombinant proteins, purification and preparation methods are listed in Table 1. The N-terminally acetylated versions of the proteins were obtained by co-expression of a plasmid encoding components of the NatB complex (Addgene), as described previously^108^. All samples were prepared in 20 mM HEPES, pH 7.4, 150 mM KCl, 10 mM MgCl_2_.

### Aggregation time course

For the aggregation reaction, aSyn was incubated at 5 mg/mL in 20 mM HEPES pH 7.4, 150 mM KCl, 10 mM MgCl_2_ in the presence of 2.5ug/mL amyloid seeds in a head-over-head rotation wheel at 37 °C for 7 days. Aliquots of 20 µL were taken each day, and Thioflavin T (ThT) was added to each aliquot at a final concentration of 5 µM. ThT fluorescence was measured at 450 nm excitation, 482 nm emission in a 386 well plate (Sarstedt) using a plate reader (CLARIOstar, BMG LABTECH).

### aSyn overexpression in Saccharomyces cerevisiae

Yeast strains used in this study are listed in Table 2. Single colonies were inoculated in appropriate synthetic dropout medium containing 2% glucose and grown over night at 30 °C under constant shaking at 180 rpm. Cells were diluted in synthetic medium containing 2% raffinose at an optical density (OD) of 0.2 at 600 nm, and grown for 12 hours at 30 °C under constant shaking at 180 rpm. To induce aSyn expression, cells were diluted in synthetic medium containing 2% galactose at a final OD of 0.2. Cells were harvested 3 h post-induction or 12 h post-induction.

**Table 2:** S. cerevisiae strains used in this study.

| Strain | Genotype | Background | Source |
| --- | --- | --- | --- |
| **αSyn-GFP** | Matalpha, aSyn-GFP:URA3, aSyn-GFP:TRP1, can1-100, his3-11.15. leu2-3,112, trp1-1, ura3-1, ade2-1 | *S. cerevisiae* W303 strain | 48 |
| **αSyn** | Matalpha, aSyn:HIS3, alphaSyn:LEU2, can1-100, his3-11.15. leu2-3,112, trp1-1, ura3-1, ade2-1 | *S. cerevisiae* W303 strain | 48 |
| **[PSI+]** | Mata, leu2-3,-112; his3-11,-15; trp1-1; ura3-1; ade1-14; can1-100, [RNQ+] | *S.cerevisiae* BY4741 strain | ATCC |

### iPSC culture and differentiation to cortical neurons

Derivation and characterization of human iPSC lines from healthy controls (SFC840-03-0, 854-03-02, SFC856-03-04), and iPSC clones from a Parkinson’s Disease patient with triplication of the SNCA gene locus (SFC831-03-01, SFC831-03-03 and SFC831-03-05) were described previously.^111,112^ iPSC were expanded in Essential 8 medium (Thermo) on Geltrex (Thermo) coated tissue culture dishes. Differentiation to cortical neurons was performed using dual-SMAD inhibition as previously described.^113^ Final plating of neural progenitors was on differentiation day 35 on polyornithine (Sigma) and Laminin521 (Thermo) coated tissue culture wells. Neurons were matured until differentiation day 56 with medium changes every 2-3 days. Then medium was removed, adherent cells were washed once with PBS and collected in PBS with a cell scraper, centrifuged at 400 g for 5 min, then the pellet was snap frozen and stored at – 80 °C until further processing. Final plating of neural progenitors was on differentiation day 35 on polyornithine (Sigma) and Laminin521 (Thermo) coated tissue culture wells. Neurons were matured until differentiation day 56 with medium changes every 2-3 days. Then medium was removed, adherent cells were washed once with PBS and collected in PBS with a cell scraper, centrifuged at 400 g for 5 min, then the pellet was snap frozen and stored at – 80 °C until further processing.

Day 56 neurons were fixed with 4% pFA and stained with a rabbit anti-aggregated alpha synuclein antibody MJFR-14-6-4-2 (abcam, ab209538, 1:20’000) and mouse anti BIII tubulin anti-body (Biolegend, 801201, 1:300).

### Wide-field fluorescence microscopy

Overexpression of GFP-tagged aSyn was induced as described above. Cells were immobilized on 8-well glass bottom dishes (ibidi) coated with concanavalin A (Sigma Aldrich) as described previously^114^. Microscopy was performed on a Deltavision microscope system with softworx 4.1.0 Softworx (Applied Precision). The system was based on an Olympus IX71 microscope, which was used with a 100×/1.4 numerical aperture (N.A.) Oil UPlanSApo objective. The images were collected with a pco.edge 5.5 camera as 1,024 × 1,024 pixel files using 1 × 1 binning and z-stacks of 0.25 µm. All images were deconvolved using standard Softworx deconvolution algorithms (enhanced ratio, high-to-medium noise filtering). Displayed images were maximum-intensity projections of at least 20 individual images. aSyn distribution rates were assessed using Fiji and its implemented Cell Counter. Cytosolic foci were distinguished based on their size (small, < 0.5 µm; large, > 0.5 µm).

### Lipid droplet staining

Overexpression of GFP tagged or untagged aSyn was induced as described above. Cells expressing untagged aSyn were stained with Nile Red as described previously^80^. BODIPY™ 493/503 (Thermo Fisher Scientific D3922) stock solution (2.5 mg/ml in DMSO) was diluted to 1 ug/ml in growth medium, mixed 1:1 with yeast culture and incubated for 10 min.

Cells were immobilized on 384-well glass bottom plates (Books, MatriPlate) coated with con-cavalin A^114^ on a temperature-controlled inverted epi-fluorescence microscope (Nikon Ti) equipped with a Spectra X LED light source and a Hamamatsu Flash 4.0 scientific complementary metal–oxide–semiconductor camera using a PlanApo 100× NA 1.4 oil-immersion objective (Nikon) and the NIS Elements software. The filters used were: Spectra emission filters 475/28 & 542/27 and DAPI/FITC/Cy3/Cy5 Quad HC Filterset with 410/504/582/669 HC Quad dichroic and a 440/521/607/700 HC quadband filter, single band emission filter GFP 510/10 and RFP 587/10 [Semrock]).

LD counts were assessed using Fiji and its implemented Cell Counter plugin. LD intensity was assessed using Fiji. Individual cells were selected as regions of interest (ROIs) using manual selection tools and the *IntDen* was measured in each ROI.

### Fluorescence recovery after photobleaching (FRAP)

Overexpression of GFP tagged aSyn was induced as described above and cells were trapped in microfluidic plates (CellASIC ONIX, Millipore). Images were acquired using a Yokogawa W1 Spinning disk on a Nikon eclipse Ti2 inverted stand microscope. The system was used with a Nikon 100x 1.45 CFI Plan Apo Oil objective. Acquired images had pixel sizes of 0.065 µm. The images were collected with an ORCA Fusion Digital CMOS camera (C14440-20UP, Hamamatsu). For imaging, we used an emission filter 525/50 and a 200mw 488nm diode laser. Images before and after photobleaching were acquired with 10 % laser intensity and 50 ms excitation time (Synuclein) or 35 % laser intensity and 150 ms excitation time (Sup35 [PSI+]). The GFP signal was bleached in a 1.36 × 1.36 µm area by using the 480-nm diode laser at 100% intensity for 200 µs each. For image analysis, we used the TrackMate Plugin from the Fiji image processing software with an estimated object diameter of 1µm. The signal from three areas was obtained for each time point: the bleached foci (I_F_), reference foci in a neighboring cell (I_R_), and an area of equal size in the background (I_B_). The fluorescence signal in the foci was normalized as follows^115^.

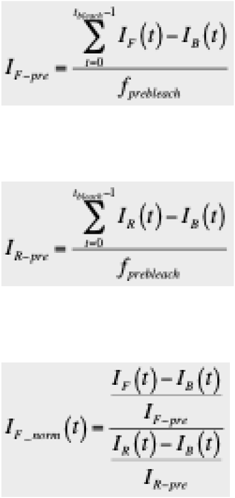

To reveal the half-time recovery, we fitted the values of I_F_norm_ with:

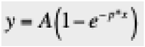

where *p* is the time constant, and *A* is the fluorescence intensity.

The half time t_1/2_ was calculated using:

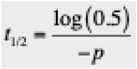

Data were analyzed, tested for statistical significance, and plotted using R software. Boxes in boxplots extend from the 25th to 75th percentiles, with a line at the median.

### Cryo-Electron tomography

Overexpression of aSyn-GFP was induced as described above. 3 µl of yeast culture were applied on glow-discharged Quantifoil^TM^ R2/1 400 mesh Au that was glow-discharged for 20 seconds. The sample was plunge-frozen in liquid ethane with a Leica EM GP2 plunger and using blotting times of 4-8 seconds.

Frozen grids were visualized with THUNDER Imager EM Cryo CLEM^TM^. The intensity of the fluorescence was enhanced to such an extent that it defined the periphery of each yeast cell and the pronounced fluorescent spots within the cytoplasm. The frozen grid samples and fluorescence grid maps of the induced cells were transferred to Aquilos 2^TM^ cryo-FIBSEM. An SEM image of the full grid was taken and overlayed with the fluorescence maps, which allowed for the identification of the cells containing aSyn-GFP foci. Yeast cells located near the center of the grid squares, which had fluorescent cytoplasmic spots, were selected for lamella preparation.

Milling was conducted according to a standard protocol, beginning with a 1 nA beam current for rough milling to produce a lamella thickness of 3 µm at a 10-degree angle, complemented by relief cuts positioned 1 µm from the edge of the lamella. The ion beam current was progressively reduced through stages—500 pA, 300 pA, 100 pA, 50 pA, and finally 30 pA—targeting thicknesses of 2.2 µm down to 150 nm, controlled by AutoTEM^TM^ software. Although the target final thickness was set to 150 nm, the final lamella were thicker (250-300 nm).

Data collection was executed using a Titan Krios G4 microscope, which is equipped with a cold-FEG, a SelectrisX energy filter was set for 20 eV zero-loss energy filtration, and a Falcon4i camera (DCI, Lausanne). Tilt-series were recorded by Tomography Software^TM^ with a total electron dose of 100 e^-^/A^2^, with a pixel size of 0.24 nm and with a defocus range of 4-5 µm. Some images were recorded (on fluorescent foci) where tilt-series recording was not possible due to the sample geometry. The tilt-series were aligned using AreTomo 1.2.5_cu11.2 ver-sion^116^, reconstructed in IMOD^117^ 4.12.18b with weighted back projection, and using a SIRT-like filter across 12 iterations. Subsequently, tomograms were denoised using the Topaz^118^ unet-3d-10a model. The projected images (Z:50-100 nm) of tomograms and recorded images were presented in the figures.

### Analysis of toxicity of aSyn overexpression

Overexpression of aSyn was induced as described above. Cells were diluted in a 2-fold series in sterile water and 5 µL was spotted onto synthetic dropout (SD-HIS-LEU) solid medium. Plates were photographed with a digital camera after 3 days of incubation at 30 °C.

### Cell lysates

Cell lysates were prepared as described previously^38,119^. In brief, *S. cerevisiae* cells were lysed using 6 cycles of glass-bead lysis in lysis buffer (20 mM HEPES pH 7.4, 150 mM KCl, 10 mM MgCl_2_, containing protease inhibitor (cOmplete™, EDTA-free Protease Inhibitor Cocktail, Roche)). Cell debris were removed by centrifugation at 10,000 g for 5 min at 4° C. Pooled iPS cell samples were lysed in lysis buffer using a tissue homogenizer (Sigma Aldrich) in four douncing cycles and cleared by two cycles of centrifugation at 10,000 g for 5 min at 4 °C. For iPS cells, wild type samples were generated by pooling neurons differentiated from 3 iPSC lines (from 3 healthy individuals); triplication samples were generated by pooling neurons differentiated from 3 iPSC lines (from a single PD individual).

For sedimentation analyses, the lysates were subjected to centrifugation at 80,000 g for 30 min at 4 °C (Beckman Coulter Optima TLX). The pellet was washed once with lysis buffer by centrifuging at 4 °C, then resuspended in lysis buffer by vortexing for 5 min and cleared by centrifugation at 8,000 g for 5 min at 4 °C.

Protein concentration was determined using a colorimetric assay (BCA Protein Assay Kit, Thermo Fischer Scientific). Biological samples were diluted to a final concentration of 2 mg/mL, purified recombinant protein samples were diluted to a final concentration of 0.2 mg/mL. All samples were kept on ice throughout the sample processing.

### Size distribution characterization

To determine molecular weight distribution, the samples were processed using a recently developed centrifugation-based filtration protocol^120^, with some minor modifications. In brief, 100 µL of sample was centrifuged at 80,000 g for 30 min at 4 °C (Beckman Coulter Optima TLX). The pellet containing the high-molecular weight fraction was washed once with lysis buffer and resuspended in 100 µL 10% (w/v) sodium deoxycholate (DOC) (Sigma Aldrich). The DOC was further diluted to 5% using lysis buffer. The supernatant was filtered through a 100-kDa MWCO filter (Microcon DNA fast flow Ultracel) at 15,300 g for 20 min at 4 °C. The flow-through containing the low molecular weight fraction was denatured by addition of 100 µL 10% (w/v) DOC. Next, 100 µL 10% (w/v) DOC was applied onto the filter membrane to denature the molecular weight fraction bound to the filter. The filter was inverted, 100 µL lysis buffer was applied on the back of the membrane, and the medium molecular weight fraction was eluted by centrifugation at 800 g for 5 min at 4 °C. Samples were frozen at -20 °C until further processing following the workflow for trypsin digestion under denaturing conditions.

### Limited proteolysis

Limited proteolysis (LiP) was carried out as described previously^38^. For pure protein samples, 10 µg of total protein was digested at an enzyme to substrate (E:S) of 1:20 and 1:100 with proteinase K from *Tritirachium album* (Sigma Aldrich). For biological samples, the E:S ratio was adjusted to match the conditions in buffer, as described previously^38^, and 100 µg of total protein was digested at an E:S of 1:50 and 1:250 with proteinase K. The samples were incubated with proteinase K for 5 min at 25 °C, and the digestion was stopped using heat inactivation at 99 °C for 3 min, followed by denaturation with 5% (w/v) DOC final concentration. Samples were frozen at -20 °C until further processing in the section on trypsin digestion under denaturing conditions.

### Trypsin digestion under denaturing conditions

All samples corresponding to a specific dataset (training dataset, validation dataset, test dataset, or samples with unknown composition) were processed together to avoid batch effects. Samples resulting from the size distribution characterization or from LiP were reduced with tris(2-carboxyethyl)phosphine (Thermo Fisher Scientific) at a final concentration of 5 mM for 30 min at 37 °C. Next, free cysteine residues were alkylated using iodoacetamide (Sigma Aldrich) at a final concentration of 40 mM for 25 min in the dark at room temperature. Samples were diluted with freshly prepared 0.1 M ammonium bicarbonate to a final concentration of 1% DOC and digested with lysyl endopeptidase LysC (Wako Chemicals) and sequencing-grade porcine trypsin (Promega), both at an enzyme/substrate ratio of 1:100, at 37 °C overnight under shaking at 800 rpm. The digestion reaction was terminated by addition of formic acid at a final concentration of 2% (v/v). Resulting peptide mixtures were loaded onto C18 96-Well Spin or 96-Well MACROSpin RPC plates (The Nest Group), desalted according to the manufacturer’s instructions, and eluted with 80% (v/v) acetonitrile. Peptides were dried in a vacuum centrifuge and solubilized in 0.1% (v/v) formic acid.

### LC-MS/MS data acquisition

Peptide digests for LiP samples were supplemented with standardized peptides for retention time correction (iRT peptides, Biognosys) and digests of ^15^N-labeled aSyn (Rpeptide) before sample acquisition. The samples were analyzed on an Orbitrap Q Exactive Plus mass spectrometer (Thermo Fisher) equipped with a nanoelectrospray ion source and a non-flow LC system (Easy-nLC 1000, Thermo Fisher). Samples corresponding to a specific dataset were acquired together in randomized order and grouped by digestion condition. Peptides were separated on a 40 cm x 0.75 µm i.d. column packed in-house with 1.9 µm C18 beads (Dr. Maisch Reprosil-Pur 120) and fractionated using a linear gradient from 5% to 35% buffer B (99.9 % acetonitrile, 0.1% FA, Carl Roth GmbH) in buffer A (0.1% FA, Carl Roth GmbH) and a flow rate of 300 nL/min. Peptides from pure protein samples were fractionated over a 90-min gradient, followed by 5 min with isocratic constant concentration of 90% buffer B; peptides from complex samples were fractionated over a 120-min gradient, followed by 5 min with isocratic constant concentration of 90% buffer B.

For shotgun LC-MS/MS data-dependent acquisition (DDA), peptides were further supplemented with yeast lysate digest. MS1 scans were acquired over a mass range of 350-1500 m/z with a resolution of 70,000. The most intense precursors that exceeded 1300 ion counts were selected for collision-induced dissociation and the corresponding MS2 spectra were acquired at a resolution of 35,000, collected for maximally 55 ms. Singly charged precursor ions and ions of undefinable charges were excluded, all multiply charged ions were used to trigger MS-MS scans followed by a dynamic exclusion for 30 s. DDA spectra were obtained for the LiP samples of the training dataset.

For data-independent acquisition (DIA) measurements, 20 variable-width DIA isolation windows were recursively acquired. The DIA isolation setup included a 1 m/z overlap between windows^38^. DIA-MS2 spectra were acquired at a resolution of 17,500 with a fixed first mass of 150 m/z and an automatic gain control (AGC) target of 1 x 10^6^. To mimic DDA fragmentation, normalized collision energy was 25, calculated based on the doubly charged center m/z of the DIA window. Injection times were automatically chosen to maximize parallelization resulting in a total duty cycle of approximately 3 s. A survey MS1 scan from 350 to 1500 m/z at a resolution of 70,000, with AGC target of 3 x 10^6^ or 120 ms injection time was acquired between the acquisitions of the full DIA isolation window sets. DIA spectra were obtained for all LiP samples.

Peptide digests for samples prepared for size characterization were supplemented with digests of ^15^N-labeled aSyn (Rpeptide) before data acquisition on a triple quadrupole/ion trap mass spectrometer (5500 QTrap, ABSciex) equipped with a nano-electrospray ion source and operated in selected reaction monitoring (SRM) mode. On-line chromatographic separation of the peptides was achieved with an Eksigent 1D-plus Nano liquid chromatography system (Eksigent/ABSciex) equipped with an 18-cm fused silica column with 75-μm inner diameter (New Objective). Columns were packed in-house with Magic C18 AQ 5-μm beads (Michrom Bioresources). Peptides were separated with a linear gradient from 1 to 35% or from 5 to 35% acetonitrile in 30 min. SRM analysis was conducted with Q1 and Q3 operated at unit resolution (0.7 *m*/*z* half maximum peak width) with a dwell time of at least 50 ms and a cycle time of less than 2 s (transition sets are described in Table S1). Collision energies were calculated according to the following formulas: CE = 0.044 *m*/*z* + 5.5 or CE = 0.055 *m*/*z* + 0.55 for doubly or triply charged precursor ions, respectively, where CE is collision energy and *m*/*z* is the mass-to-charge ratio of the precursor ion.

### Peptide and protein identification

DDA spectra of the training dataset were analyzed with Spectronaut (Biognosys AG, version 13) and the built-in search engine Pulsar with default settings with the following modifications: Digestion enzyme specificity was set to semi-tryptic with Trypsin/P and LysC. Labeling was applied, and labels in channel 2 were specified as 15N(1), 15N(2), 15N(3), and 15N(4). Search criteria included carbamidomethylation of cysteine as a fixed modification, as well as oxidation of methionine and acetylation (protein N-terminus) as variable modifications. Up to two missed cleavages were allowed. The DDA files were searched against the *S. cerevisiae* (strain S288c) Uniprot fasta database (version October 2017), the aSyn sequence, the Biognosys’ iRT peptides fasta database, and a contaminant fasta database (described in Malinovska *et al*, 2022^38^). The libraries were generated using the library generation functionality of Pulsar with default settings. For the compositional predictions, DIA spectra were analyzed with Spectronaut (Biognosys AG, version 13) using the default settings, and the spectral libraries generated as described above. The dynamic mass tolerance strategy was applied to calculate the ideal mass tolerances for data extraction, and no correction factor was applied (correction factor = 1). The local (non-linear) regression method was used for iRT calibration using the iRT kit peptides. The mutated decoy method was used to generate label-free decoys. The false discovery rate (FDR) was estimated with the mProphet approach and was set to 1% at both the peptide precursor and protein levels. Precursor ion quantities for aSyn were extracted. For the analysis of global structural changes, DIA spectra were analyzed with Spectronaut (Biognosys AG, version 15) using the library-free workflow directDIA. The search was performed using the Biognosys’ iRT peptides fasta database and a contaminant fasta database as well as the *S. cerevisiae* (strain S288c) Uniprot fasta database (version October 2017) and the aSyn sequence for the *S. cerevisiae* overexpression dataset or using the human Uniprot fasta database (version October 2017) for patient iPSC-derived neurons. The search was performed with default settings, with minor modifications. Digestion enzyme specificity was set to semi-tryptic with Trypsin/P and LysC. Search criteria included carbamidomethylation of cysteine as a fixed modification as well as oxidation of methionine and acetylation (protein N-terminus) as variable modifications. Up to two missed cleavages were allowed.

## Data Analysis

All statistical analyses were performed with the statistical language R (v. 4.0.4)^121^. Unless stated otherwise, functions from the stats package or the tidyverse package^122^ were used.

### Data processing

Run outliers due to measurements errors were identified based on the intensity distributions of the ^15^N-labeled aSyn peptides. Runs with a significant difference in intensity distributions (Wilcoxon test, p-value < 0.0001) compared to all other technical replicates originating from the same source were excluded. Precursor peptides with intensities below the detection threshold were removed as were precursors identified in less than 75% of the replicates. Peptide quantities were calculated as sum of all precursors. Within each dataset, run intensities were normalized across all runs in both digestion conditions. Run outliers due to sample processing were identified by Spearman correlation to Huber M-estimator, and runs with R^2^ < 0.8 were excluded. The median coefficient of variation per peptide in each digestion condition of the training dataset was determined, and peptides with outliers (median CV > Q3 + 1.5 IQR) were removed.

Missing values were imputed based on their missing type: values missing at random (MAR) were defined as peptides where one replicate had a missing value; values missing not at random (MNAR) were defined as peptides where no values were detected in all four replicates. For MAR values, the imputed value was sampled from a normal distribution with the mean value of that peptide and the standard deviation of that peptide. For MNAR values, imputed values were calculated as the threshold of detection plus an error sampled from a normal distribution with mean 0 and the global standard deviation within the dataset.

The structural information of the digestion pattern is not determined by the absolute abundance of a peptide, but by the relative intensity of each peptide of the protein, rendering the fingerprint independent from the total protein concentration. Thus, the data qualifies as compositional data as the key principle of compositional data is scale invariance^123^. The treatment of mass spectrometry data as compositional data has been also suggested previously, but in a different context^124^. Therefore, each dataset was treated as compositional dataset. It was first log_2_-transformed to achieve normal distribution. Then *closure* was created: The peptide intensities were divided by the sum of intensities in each sample, yielding proportional data.

### Characterization of structural fingerprints

The discriminatory power of the structural fingerprints was assessed using a Kruskal-Wallis H-test as well as a linear mixed effects model as described previously^125^. Feature values were hierarchically clustered in columns according to Pearson’s correlation coefficients using Ward’s agglomerative clustering method (Ward.D2)^126^. Data-dimensionality reduction was achieved using principal component analysis (PCA), linear discrimiant analysis (LDA; with the *lda* function of the MASS package^127^) and t-distributed stochastic neighbor embedding with the *tsne* function of the M3C package^128^ and perplex value of 20.

We attempted to generate a minimal fingerprint comprising the most distinctive features, but it did not outperform the full fingerprint (see below).

### Composition prediction

The composition prediction was based on a multivariate mixing model, where the composition of a sample comprises a mixture of end members, which are defined by the structural fingerprints of the pure structures. This resulted in an overdetermined inverse linear problem of the form Ax = b (*i*), with the matrix A containing the first to n^th^ feature value of each structure in the training dataset (𝑚 = disordered monomer, 𝐴 = oligomer type-A*, ℎ = lipid-bound helical, 𝐵= oligomer type-B*,𝑓 = amyloid fibril, asterisks in equations (i-iv) below annotate non-acetylated variants); the vector x containing the contributions of each end member to the composition; and the vector b containing the first to n^th^ feature value of the sample with unknown contribution 𝑢. The feature values 𝑣 were calculated as the mean value of a given structure or input sample.

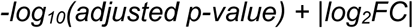

This equation can be solved using constrained least squares regression with the lsei algorithm^129^ implemented in the limSolve package^130^. We determined the detected features for each dataset and excluded features missing globally in the sample dataset. For the validation dataset, we determined the globally missing features in a background-dependent manner.

The equality constraint (*ii*) was set by the requirement of the contributions 𝑥_𝑚−𝑓_ to equal 1.

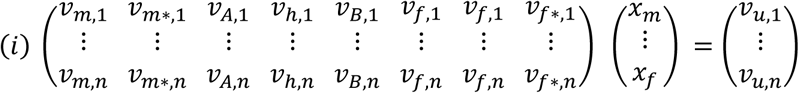

The inequality constrains limited the contributions 𝑥_𝑚−𝑓_ to be positive numbers (*iii*).

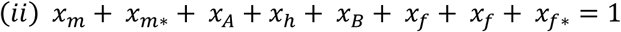

The size distribution of the different *in vitro* structural conformations can be used as basic discriminatory measure (Figure S6). Thus, we used the determined size distributions of each sample as additional inequality constraints, allowing for 10% error in the sample determination, where 𝑙𝑚𝑤 is the ratio of the low molecular weight species, 𝑚𝑚𝑤 is the ratio of the medium molecular weight species, and ℎ𝑚𝑤 is the ratio of the high molecular weight species (*iv*). The size distributions in each data set are shown in Figure S6.

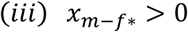

In addition, we used the coefficient of variation (CV) to calculate the weighting coefficients vector; we calculated the median CV for each feature in the training dataset and inversely scaled the median CV values to range from 1 for the lowest median CV to 0 for the highest median CV.

To account for the variability in the fingerprint that is introduced through sample handling and/or measurement precision, we applied the Bayesian Compositional Estimator (BCE) algorithm^42^. This algorithm incorporates prior probability distributions of the fingerprint values as well as the unknown sample values. We used the resulting vector containing the solution of the least squares problem as starting point for the Bayesian composition estimator *bce1* implemented in the BCE package^42^. To determine the prior probability distribution of the input matrix A, we defined the boundaries for the sampling of the matrix A as the mean of the feature value ± one standard deviation. The BCE algorithm also allows the assessment of posterior probability distributions through Markov chain Monte Carlo random walk sampling^42^. For this, we used a jump length between 0.5×10^-6^ and 2×10^-6^ for the input A and between 1×10^-4^ and 4×10^-4^ for the composition vector x and performed 10^6^ iterations. We confirmed convergence of the Markov chain Monte Carlo procedure by visual inspection of the residuals^42^. Analogous to the *lsei* function, we used weighting coefficients based on the coefficient of variation, both for the input matrix A and the sample vector b, calculated as described above. The contributions from the different disordered monomer and amyloid fibril species were summed up to determine the ratios of the structures.

### Selection criteria for fingerprints

Although peptide identification was substantially improved using data-independent acquisition (DIA) methods^7,8^ and yielded a more consistent data set across samples, we observed a certain fluctuation in the recovery efficiency of the peptides. In particular, we observed a striking difference in the recovery efficiency for samples in buffer compared to samples in complex background.

However, since compositional data shows subcompositional coherence^123^, we used subsets of the features and removed the non-identified peptides for the predictions in each dataset, to generate the full fingerprint.

To account for the stochastic aspect of the mass spectrometry methods, we also generated a robust fingerprint, and chose the peptides that were identified in at least 10 out of the 11 data sets used. This corresponds to a recovery rate of 84%, with a total of 69 peptides, of which 58 were identified in all data sets and 11 were missing in only one. We generally observe a similar recovery rate when re-acquiring samples.

Selecting non-redundant features can improve the prediction outcome by preventing overfitting, therefore we tested feature selection by stepwise elimination method, minimizing Willk’s lambda^131^. A minimal fingerprint was generated by minimizing Wilks’ Lambda using the *greedy.wilks* function of the klaR package^132^ with a niveau value of 0.03. However, the features showed a collinear relationship, despite data transformation. This could result from the fact that the intensities of the peptides are not independent – intensities of peptides resulting from proteinase K cleavage increase while the intensity of the non-cleaved peptide decreases. Other feature selection methods might prove more effective.

### Error determination

The prediction precision was assessed using two measures. The first was the value of the minimized quadratic function at the solution of the least squares regression, as computed by the *lsei* function. Since this value is dependent on the number of features included in the model, we calculated the relative sum of squared residuals by scaling the squared sum of residuals (SSR), using the smallest SSR in the cross-validation prediction as minimum and smallest SSR in the prediction with random values as maximum. This allowed comparisons of the prediction performance across different datasets. As the second measure, we analyzed the squared residuals for each feature and the differences in their distributions.

The prediction error was determined in the jackknife cross-validation, the validation dataset, and the test dataset by using the sum of weighted last squares between the observed and the expected compositions. Errors in the expected structures were weighted by factor 2, and the remaining deviations had a weight value of 1.

### Size characterization

SRM measurements for size characterizations were analyzed using Skyline (MacCoss Lab Software), peptide ratio results were exported and processed with the statistical language R (v. 4.0.4)^121^. The relative protein abundances in the size fractions were calculated as the mean ratio to standard. The size ratios were determined as proportions of the protein abundances in the three size fractions of a sample.

### Proteolytic resistance analysis

The proteolytic resistance was assessed in the strong digestion conditions (E:S 1:20) using fully-tryptic peptides with no missed cleavages. It was calculated as the ratio of the intensity of fully-tryptic peptides in the LiP condition to the Trypsin-only condition, where the sample has not been treated with PK. In cases where the peptide intensities in both conditions did not show statistical evidence for a difference (student t-test, p>0.05), the proteolytic resistance was set to 1.

### Quasi-residue-level analysis of structural differences

We explored the possibilities of using the full structural fingerprints for the analysis of structural features. As the structural fingerprints can only be informative when comparing two or more conditions, we compared each structure in the library with the other structures (e.g. Monomer vs Fibril, Monomer vs helical lipid-bound, Monomer vs. Oligomer A*, etc). This comparison allows us to pinpoint peptides that are different in the two structures. Since we have a high coverage of the protein, we obtain multiple peptides covering one region. In order to condense the information per amino acid position, we performed two orthogonal analyses.

In the first approach, we assessed the distribution of different and non-different peptides at each position. For this, we assigned the peptides into two categories, those with strong evidence for a difference (q-value < 0.05, “different”) and those which do not pass the significance threshold (“non-different”). Then, we assessed whether the distribution of categories along the sequence was random using a chi-squared test for independence. Since this test is more robust for category sizes >5, and we do not always have 5 or more peptides covering each residue of the sequence, we also ran the chi-squared test on binned data, where we pooled information across the median length of a peptide (13 amino acids). This approach reduces the resolution but allows for robust statistical analysis of the distributions.

In the second approach, we calculated a dissimilarity score for each position, based on the significance (q-value) and magnitude (log2(FC)) of the change detected in all peptides covering that position. The dissimilarity score was calculated per amino acid position as: median (-log10(q-value) * |log2FC|), and it was scaled to range from 0 to 1. Both fully tryptic and half tryptic peptides were included in the analysis. A high score value indicates a higher probability for a structural alteration at a given position. We can define a threshold value for the dissimilarity score based on pre-defined cutoffs for the significance (q-value < 0.05) and the magnitude (|log2FC|>1). Alternatively, we established an independent threshold value based on the precision of locating altered peptides in the vicinity of the active site upon binding of the vitamin-D-binding protein GC (PDB 1J78) to its known ligand vitamin D^133^. To do this, we iterated through the dissimilarity scores (ranging from 0 to 1) for the test protein with and without added ligand. At each score, we selected regions exceeding that dissimilarity value and calculated precision using the formula TP / (TP + FP), where TP represents true positives (regions located within 6.4 Å of the active site) and FP represents regions located farther from the active site (> 6.4 Å). The significance threshold was chosen as the point with the highest precision at the highest cutoff. We also checked the scores and threshold values against other test proteins. Overall, we observed consistent results in our comparisons of different structural forms of aSyn in all of these analyses (i.e., using the dissimilarity score with both threshold values or using the difference assessment (Fig. S2).

### Global alterations

To identify biological pathways and compartments affected by aSyn overexpression, a protein set enrichment analysis was performed using the data obtained at mild digestion conditions. This approach was based on the gene set enrichment analysis reported previously^134,135^. Compared to a classic Gene Ontology enrichment analysis, the protein set enrichment analysis defines whether a defined set of proteins shows statistically significant differences between two states, without applying any arbitrary significance threshold. This unbiased approach has the advantage of overcoming the possible loss of statistical power due to thresholding.

The following rank metric score was computed for every detected LiP peptide:

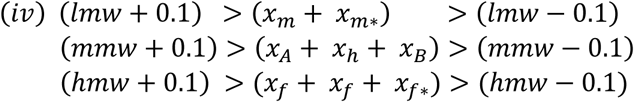

where adjusted p-value and log_2_FC result from a differential abundance testing, using unpaired two-sample Student’s t-test, performed at peptide level with the software Spectronaut (Biognosys AG, version 13). The best scoring peptide per protein (“sentinel peptide”) was selected. The final list of sentinel peptides was ranked and sorted by descending numerical order. The ranked list was analyzed using the *prerank* function of the GSEAPY python package (version 0.10.8). Protein sets were defined using the “Cellular component” terms in the Gene Ontology (GO) database^59^ and the full list of *Saccharomyces cerevisiae* metabolic pathways from the Kyoto Encyclopedia of Genes and Genomes (KEGG) PATHWAY Database^58^. For GO sets, GO ontologies were extracted from the <u>go-basic.obo</u> file (2022-03-22 release). Terms with more than 500 or less than 5 annotated genes were excluded from the analysis. Similarly, KEGG pathways with more than 100 or less than 3 annotated genes were excluded from the enrichment analysis. The number of permutations for significance computations was set to 5000 and maximum and minimum number of proteins from proteins sets in the ranked list was set to 500 and 5, respectively. As result of the protein set enrichment analysis, an enrichment score (ES) is calculated. The ES score was calculated by walking down the ranked list of LiP sentinel peptides and increasing a running-sum statistic whenever the associated protein is present in the defined protein set and decreasing it when peptides are not associated to any protein in the defined protein set. The final enrichment score is where the maximum deviation from zero is encountered by walking down the list of LiP sentinels. The ES score was calculated by walking down the ranked list of LiP sentinel peptides and increasing a running-sum statistic whenever the associated protein is present in the defined protein set and decreasing it when peptides are not associated to any protein in the defined protein set. The final enrichment score is where the maximum deviation from zero is encountered by walking down the list of LiP sentinels. The ES was finally normalized by dividing the ES by the mean of the permutation enrichment values and false discovery rate (FDR) was calculated using permutation testing by generating a null distribution of normalized enriched values (NES). To select for peptides which are overrepresented at the top of the ranked LiP list, we filtered the output for ES > 0 and FDR < 0.25. In order to visualize the result of the protein set enrichment analysis, we plotted NES scores for significantly enriched protein sets (ES > 0 and FDR < 0.25). For high scoring protein sets we reported the leading-edge proteins, meaning the set of proteins for which a LiP peptide is found in the ranked list with a score higher than the ES value. Global structural alterations were analyzed as described previously using the MSstatsLiP package^38^.

## Data visualization

The data were visualized using the statistical language R (v. 4.0.4)^121^ and the packages ggplot2^136^, ggpubr ^137^, ggsci ^138^, and gghighlight ^139^.

