## Supplementary material for "Deconvolving the structural heterogeneity of alpha-Synuclein in *vitro* and *in situ*": Malinovska et al Supplement

### Figure S1

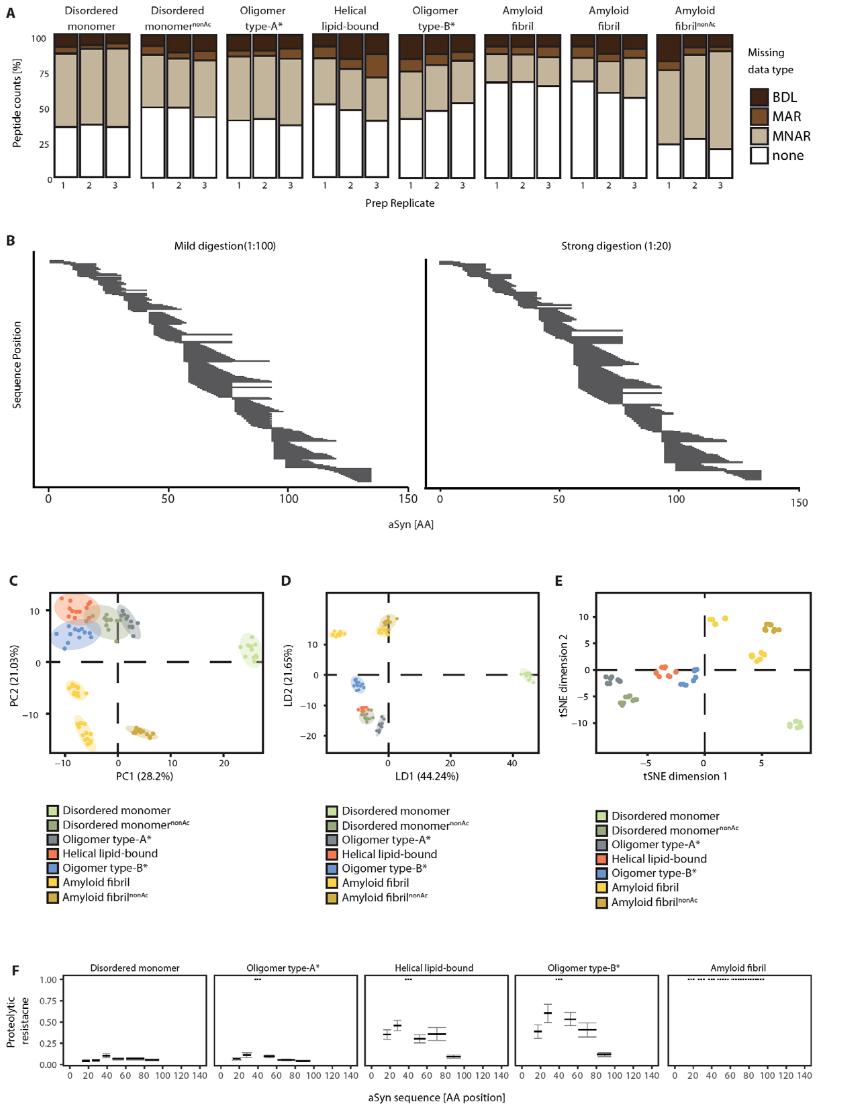


**Figure S1: Characterization of the structure-specific digestion patterns. (A)** Completeness of the peptides identified using data-independent acquisition (DIA) in the conformational samples for both digestion conditions. Independent preparations are displayed nextto each other. Missing types are displayed as fraction of total (in [%]). Peptides identified in all four technical replicates in each structureare classified as not missing (none, white); peptides missing in one or more conformations are classified as not missing at random (MNAR, light brown); peptides missing or below the intensity threshold in one of four technical replicates of a structure are classifiedmissing at random (MAR, brown); peptides with intensities below the threshold in more than one replicate are classified below detection limit (BDL, dark brown). **(B)** Peptides from mild (left) and strong (right) digestion are ranked according to their location along the amino acid sequence and mapped to the sequence of aSyn. **(C-E)** Distribution of structural fingerprints upon dimensionality reduction using **C)** Principal component analysis, **D)** linear discriminant analysis, and **E)** t-distributed stochastic neighbor embedding (tSNE) analysis. Groups are colored by structure. **(F)** Proteolytic resistances of the six fully-tryptic peptides from aSyn reference structures are plotted along the sequence of aSyn. Peptides with no statistical evidence for a difference in intensity (student t-test, p>0.05) are visualized using a dotted line. Error bars represent standard error of the mean (SEM).

**Figure S2**

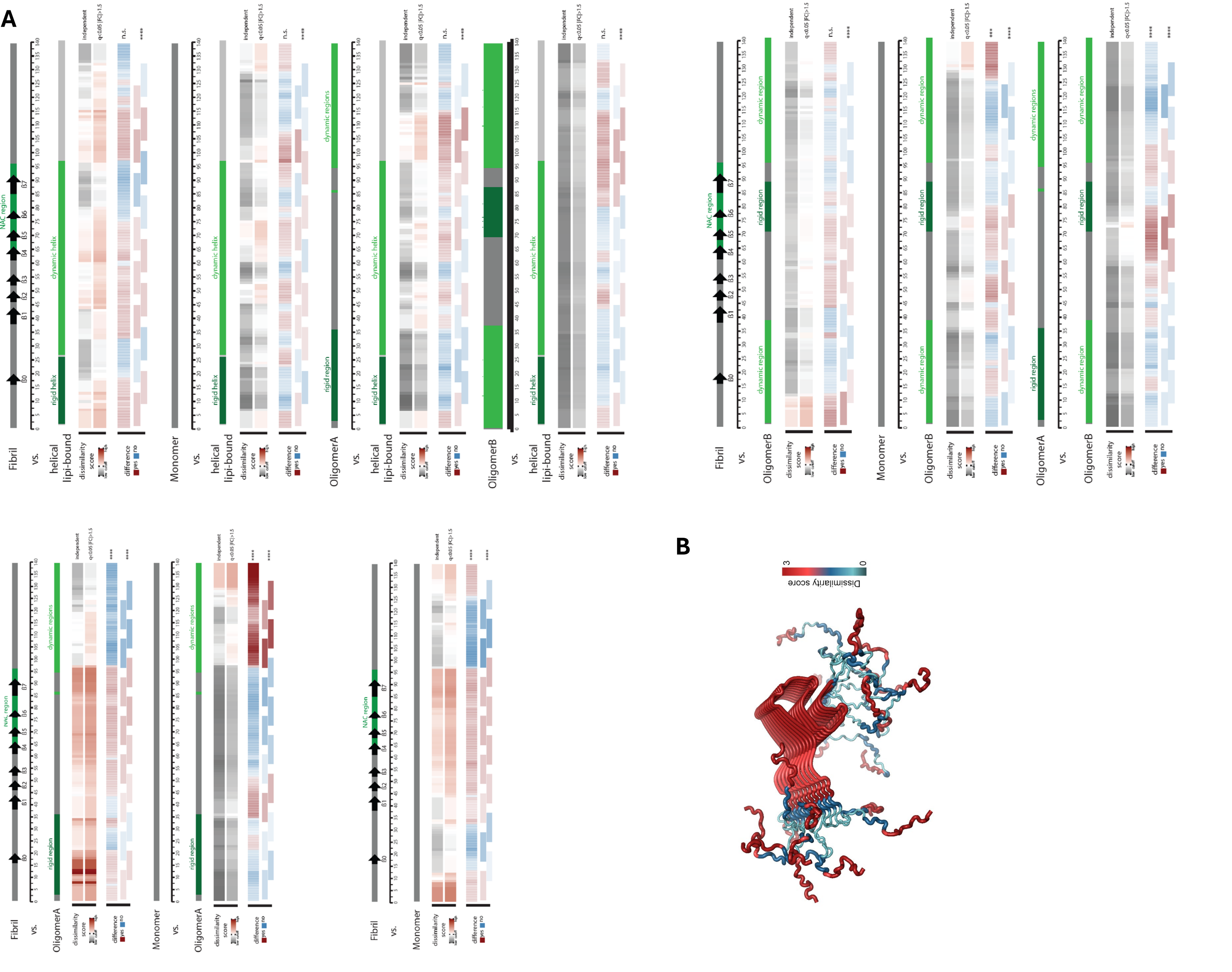

**Figure S2: (A) Pairwise comparison of digestion patterns of the reference aSyn structures.** Structural features of the two conformations being compared are visualized along the sequence of aSyn. The dissimilarity score is calculated as the median of the product of –log10(q-value) and |log2(fold change)| and scaled to range between 0 and 1. Thresholds for the dissimilarity are either determined based on the precision of locating altered peptides in the vicinity of the active side upon ligand binding (upper) or based pre-defined cutoffs for the significance (q-value < 0.05) and the magnitude (|FC|>1.5) (lower). The randomness of distribution of difference categories (yes, dark red; no, blue) on amino acid level (upper) or spanning a region of 13amino acids (lower) is assessed using a chi-squared test for independence (****: p-value < 0.0001; ***: p-value < 0.001; **: p-value < 0.01; *: p-value < 0.05; n.s.: p-value > 0.05). **(B) Comparison of in vitro aSyn monomeric and fibril structures.** Structural dissimilarity scores calculated from LiP-MS data from in vitro aSyn monomer and fibril were mapped onto fibril polymorph 2b structure (PDB id: 2N0A). The dissimilarity score is the median of the product of –log10(q-value) and |log2(fold change)|, scaled to range between 0 and 1. The higher the score, the larger the differences between the two structures.

### Figure S3

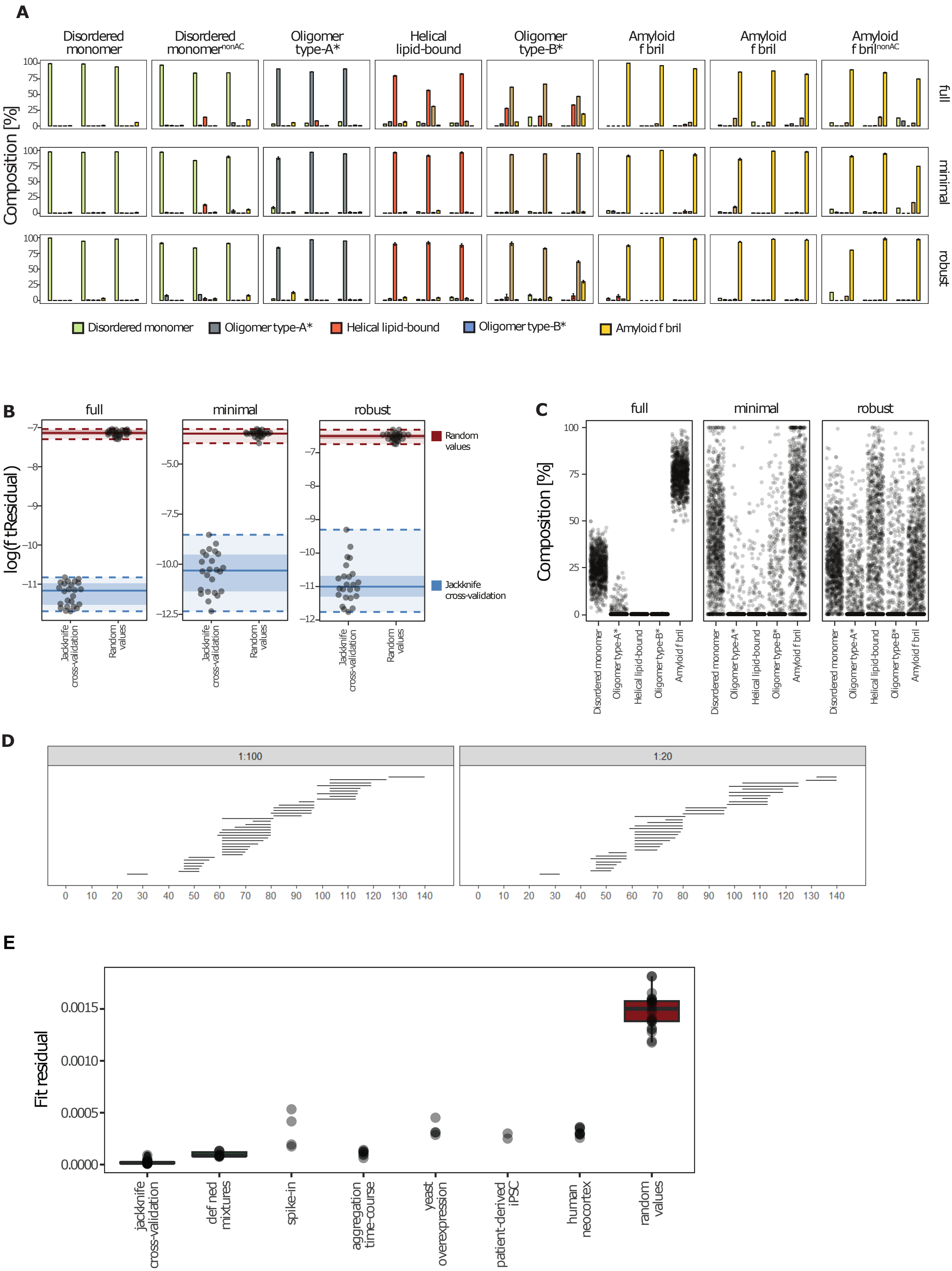


**Figure S3: A prediction model to determine conformational composition. (A)** Predictions of sample compositions using jackknife cross-validation with the full (upper row), minimal (middle row) and robust (lower row) fingerprint. The results of the three preparations for each sample are displayed next to each other. The contribution of each structure is plotted as the fraction of the total (in [%]), and the identities of the predicted structures are indicated by color, as indicated. Errors are displayed as ranges and represent 95% confidence intervals of the Bayesian resampling. **(B)** Log transformed fit residuals for the jackknife cross-validations in A and for predictions using random values. Ranges are visualized by color (blue, jackknife cross-validation; red, random values; thick lines, median; dotted lines, min and max; dark shade, IQR). **(C)** Cumulated conformation composition predictions using random values and the full, minimal or robust fingerprint. Each point represents one simulation of 1000 iterations. **(D)** The plots show aSyn peptides within the robust subset of the structural library dataset (i.e., the robust fingerprint). Each line indicates an individual peptide, mapped along the sequence of aSyn. The fingerprint is shown for the two indicated digestion conditions (PK:substrate) **(E)** Scatterplot of fit residuals in jackknife cross-validation, predictions in test and validation data sets (defined mixtures and spike-in), predictions in samples with unknown compositions (aggregation time-course, aSyn overexpression in S. cerevisiae, patient iPSC-derived neurons and human cortex), and predictions with random values. The distribution of fit residuals in data sets with >7 samples are also visualized using boxplots.

### Figure S4

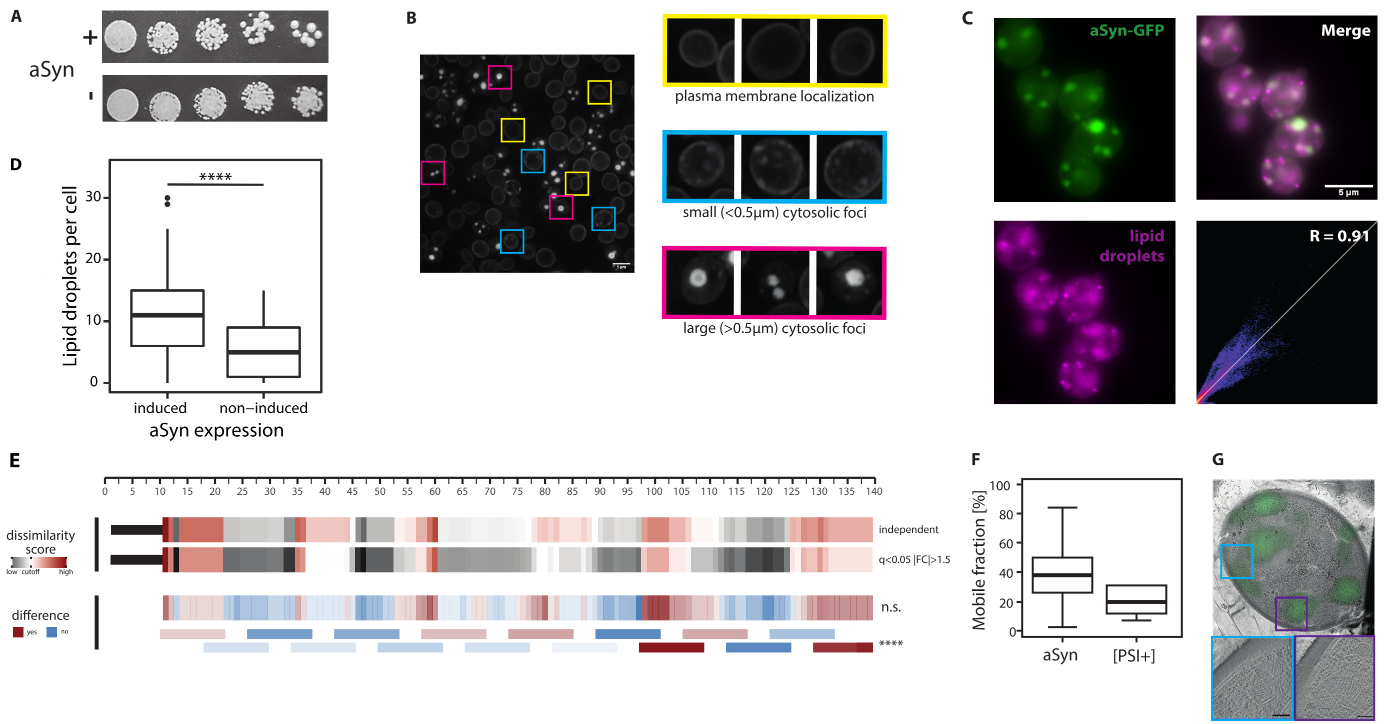


**Figure S4: Effects of aSyn overexpression in S. cerevisiae (A)** Assessment of toxicity caused by aSyn expression. Photographs of two-fold serial dilution for analysis of growth of *S. cerevisiae* that expresses aSyn (+) or not (-). **(B)** Assessment of aSyn distribution 12 hours post-induction. Yeast cells are manually assigned to three categories: plasma membrane localization (yellow), small (< 0.5 µm) cytosolic foci (blue) or large (> 0.5 µm) cytosolic foci. Three examples of each category are shown. **(C)** Localization of aSyn-GFP and lipid droplets (stained with Nile Red) 12 hours post-induction. Correlation between the two markers is shown. **(D)** Number of LDs with and without 12h induction of aSyn expression in *S. cerevisiae* expressing wild type aSyn LDs counted in a maximum projection of 25 z-stacks, visualized by BODIPY staining. **(E)** Comparison of the structure-specific digestion pattern of lysates of *S. cerevisiae* that express GFP-tagged aSyn at 3 h post-induction and 12 h post-induction. The dissimilarity score is calculated though multiplication of the q-value and |log2(fold change)| and scaled to range between 0 and 1. Thresholds for the dissimilarity are either determined based on the precision of locating altered peptides in the vicinity of the active site upon ligand binding (upper) or based pre-defined cutoffs for the significance (q-value < 0.05) and the magnitude (|FC|>1.5) (lower). The randomness of distribution of difference categories (yes, dark red; no, blue) on amino acid level (upper) or spanning a region of 13amino acids (lower) is assessed using a chi-squared test for independence (****: p-value < 0.0001; ***: p-value < 0.001; **: p-value < 0.01; *: p-value < 0.05; n.s.: p-value > 0.05). **(F)** Quantification of the mobile fraction in S. cerevisiae expressing GFP-tagged aSyn at 12 h post-induction (left, n = 37 foci) or GFP-tagged [PSI+] (right, n = 27 foci). **(G)** Electron tomograms of *S. cerevisiae* expressing GFP-tagged aSyn, overlayed with fluorescence signal for identification of GFP signal. Two areas with GFP-positive (continues on next page)

#### Figure S5

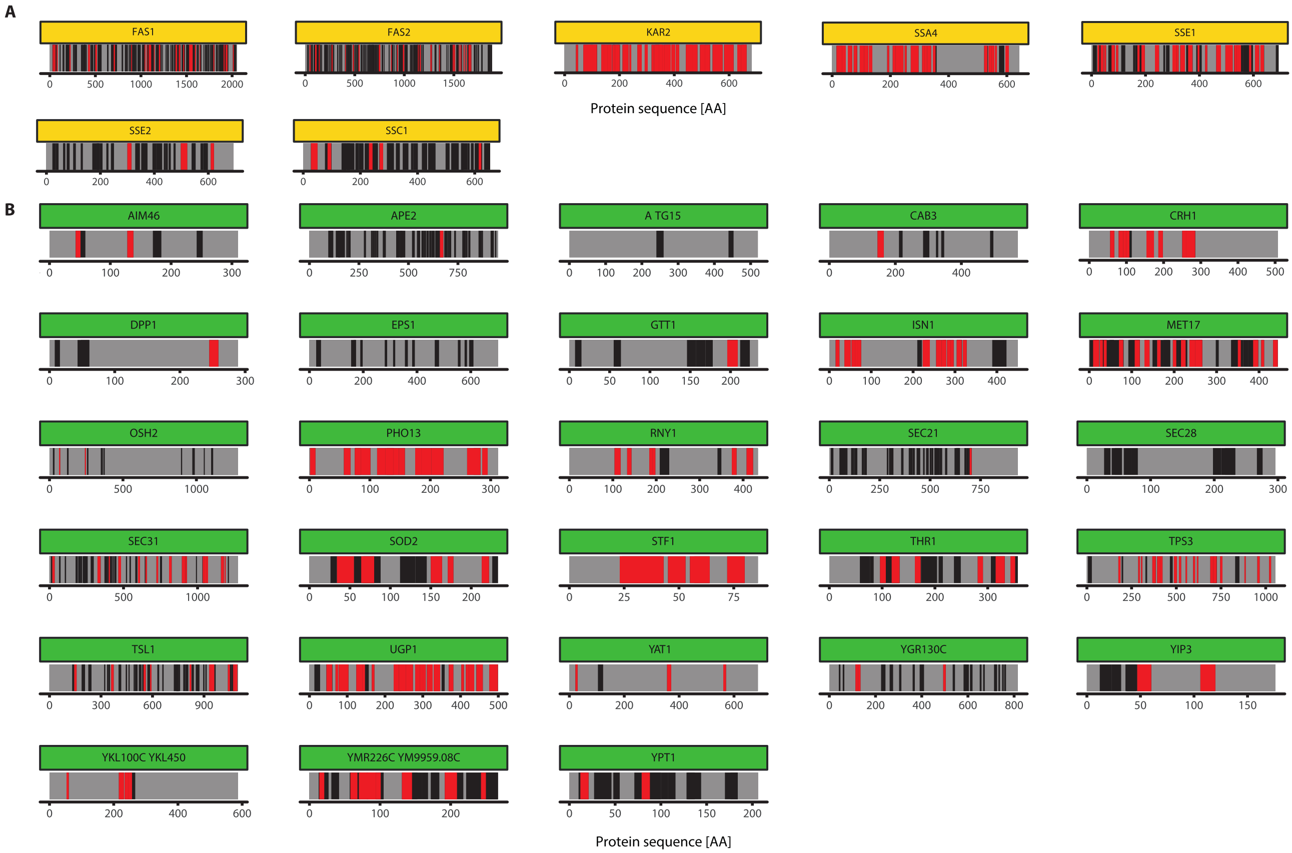


**Figure S5: visualization of structural alterations in aSyn interactors (A, yellow) and modulators of toxicity (B, green).** Peptides displaying a significant change in the mild digestion condition are plotted along the sequence of the protein and highlighted in red. Peptide with no significant change are shown in black, positions not identified are shown in grey.

### Figure S6

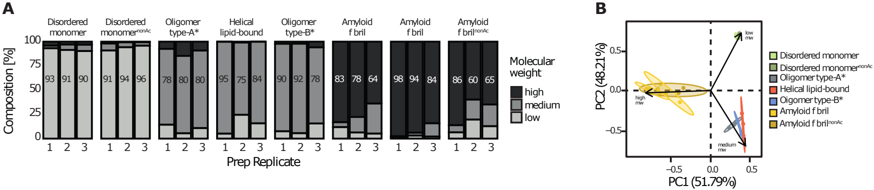


**Figure S6: Size characterization of aSyn in vitro structures. (A)** Size distributions in the in vitro samples analysed to generate our prediction algorithm (see Fig 1A). Three independent preparations of each structure were analyzed with each visualized in a bar chart. The molecular weight distribution is presented as fraction of the total (in [%]), with high molecular weight in black, medium molecular weight in dark gray, and low molecular weight in light gray. **(B)** Principal component analysis of the weight distributions in the conformational samples. Arrows in the biplot represent the variables, groups are colored by structure as indicated (non-acetylated versions of the disordered monomer and the amyloid fibril are indicated by darker shade of the group color).
